# Sporadic activation of an oxidative stress-dependent NRF2–p53 signaling network in breast epithelial spheroids and premalignancies

**DOI:** 10.1101/862474

**Authors:** Elizabeth J. Pereira, Joseph S. Burns, Christina Y. Lee, Taylor Marohl, Delia Calderon, Lixin Wang, Kristen A. Atkins, Chun-Chao Wang, Kevin A. Janes

## Abstract

Breast–mammary epithelial cells experience different local environments during tissue development and tumorigenesis. Microenvironmental heterogeneity gives rise to distinct cell-regulatory states whose identity and importance are just beginning to be appreciated. Cellular states diversify when clonal 3D spheroids are cultured in basement membrane, and prior transcriptomic analyses identified a state associated with stress tolerance and poor response to anticancer therapeutics. Here, we examined the regulation of this state and found that it is jointly coordinated by the NRF2 and p53 pathways, which are co-stabilized by spontaneous oxidative stress within the 3D cultures. Inhibition of NRF2 or p53 individually disrupts some of the transcripts defining the regulatory state but does not yield a notable phenotype in nontransformed breast epithelial cells. In contrast, combined perturbation prevents 3D growth in an oxidative stress-dependent manner. By integrating systems models of NRF2 and p53 signaling together as a single oxidative-stress network, we recapitulate these observations and make predictions about oxidative stress profiles during 3D growth. Similar coordination of NRF2 and p53 signaling is observed in normal breast epithelial tissue and hormone-negative ductal carcinoma in situ lesions. However, the pathways are uncoupled in triple-negative breast cancer, a subtype in which p53 is usually mutated. Using the integrated model, we reconcile the different NRF2-knockdown phenotypes of triple-negative cancer lines with their inferred handling of oxidative stress. Our results point to an oxidative stress-tolerance network that is important for single cells during glandular development and the early stages of breast cancer.

**One Sentence Summary:** Reactive oxygen species co-stabilize a non-oncogene and a tumor suppressor for triple-negative breast cancer when cells are surrounded by basement-membrane ECM.

## Introduction

Among glandular tissues, the breast–mammary epithelium is unique because of the dramatic expansion and reorganization that occurs after birth (*1*). During puberty, a branched network of epithelial ducts is pioneered by terminal end buds (TEBs), which emerge from the rudimentary gland and extend into the surrounding mesenchyme (*2*). TEBs contain a mixture of proliferating stem–progenitor cells and differentiating cells fated to the secretory luminal-epithelial or contractile basal-myoepithelial lineages. During morphogenesis, TEB cells are dynamically exposed to different microenvironments that inform final organization of the gland (*3*). Some microenvironmental cues are supportive or instructive to cells [hormones (*4*), growth factors (*5*), basement membrane (*6*)]. Others are deleterious or lethal [loss of polarity (*7*), detachment (*8*), ER stress (*9*)]. All of these cues are reconfigured aberrantly and heterogeneously during the early stages of breast–mammary cancer (*10–12*).

Stress and survival signals also juxtapose when breast–mammary epithelial cells are grown in 3D culture with reconstituted basement membrane ECM (*13, 14*). Combining the appropriate adhesive and soluble cues yields TEB-like behavior in 3D-cultured multicellular epithelial fragments from the mammary gland (*7*). For single-cell cultures that reliably organize as 3D structures, clones or progenitors must iteratively proliferate, maintain cell-cell adhesions, and coordinate function to establish a multicellular ecosystem (*15, 16*). Cell-regulatory states diversify within 3D organoids of primary breast–mammary epithelia (*17–19*) and also in the simplest 3D spheroids of isogenic cell lines (*20–23*). Identifying such cell-regulatory heterogeneities is important, because there are parallels to in situ lesions of the breast, where premalignant cells must survive and proliferate in the duct (*24, 25*).

Previously, we identified a cluster of transcripts (Fig. 1A) that covaries heterogeneously among hormone-negative, basal breast epithelial cells grown as 3D spheroids (*24*). The cluster contains *KRT5* (a PAM50 classifier for basal-like breast cancer) (*26*) along with multiple stress-tolerance genes, including *JUND* (*27*), *CDKN1A* (*28*), *MUS81* (*29*), and *HSPE1* (*30*). The transcripts in this cluster were among the strongest and most-negative predictors of breast-cancer response to chemotherapy and targeted agents in an independent clinical trial (*31*). We reported that individual genes in the cluster have complex time- and microenvironment-dependent relationships in 3D spheroids, animal models of ductal carcinoma in situ (DCIS), and clinical hormone-negative premalignancies (*24*). However, the overarching regulation of the cluster was not determined.

**Fig. 1.**
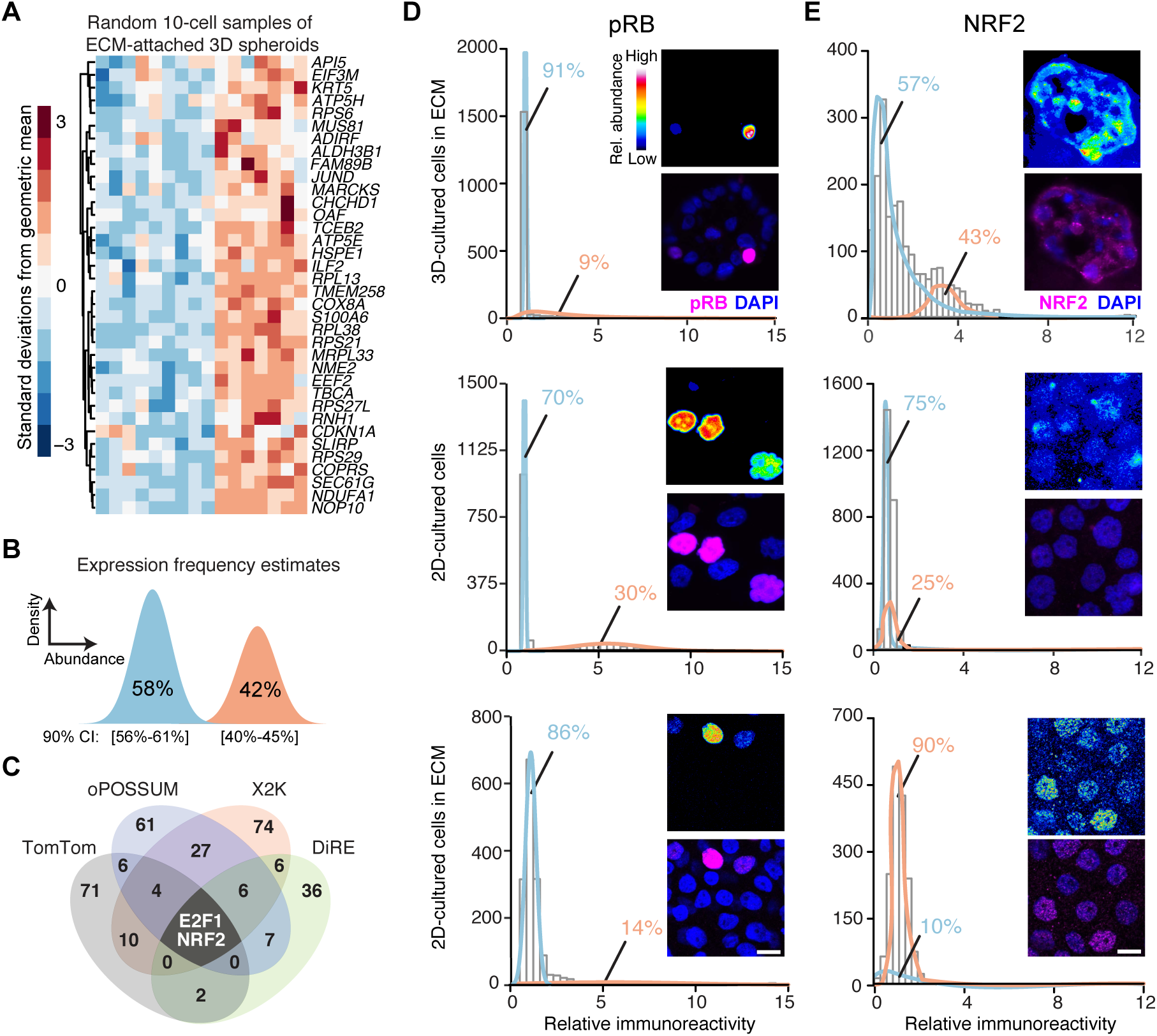
Transcriptomic fluctuations of ECM-cultured breast epithelial spheroids reveal a gene cluster associated with heterogeneous NRF2 stabilization in a 3D-specific environment. (**A**) Microarray profiles of ECM-attached basal-like MCF10A-5E breast epithelial cells randomly collected as 10-cell pools (*n* = 16) from 3D-cultured spheroids after 10 days (*20*). (**B**) Maximum-likelihood inference (*38*) parameterizes a two-state distribution of transcript abundances for the gene cluster in (A). Inferred expression frequencies are shown as the maximum likelihood estimate with 90% confidence interval (CI). (**C**) Promoter-bioinformatics methods converge upon NRF2 and E2F1 as candidate regulators of the gene cluster. (**D** and **E**) Quantitative immunofluorescence of (D) hyperphosphorylated RB (pRB, an upstream proxy of active E2F1) and (E) NRF2 in 3D culture with ECM (upper), 2D culture (middle), and 2D culture with ECM (lower). Expression frequencies for a two-state lognormal mixture model (preferred over a one-state model by *F* test; *p* < 0.05) were calculated by nonlinear least squares of 60 histogram bins collected from *n* = 1100–1600 of cells quantified from 100-200 spheroids from two separate 3D cultures. For each subpanel, representative pseudocolored images are shown in the upper right inset and merged (magenta) with DAPI nuclear counterstain (blue) in the lower right inset. Scale bar is 10 µm.

Here, we find that regulatory-state heterogeneity emerges from the coordinated action of two stress-responsive transcription factors—NRF2 (*32, 33*) and p53 (*34*)—which become stabilized posttranslationally when breast-epithelial cells variably experience oxidative stress in 3D culture. Genetic disruption of NRF2 signaling alters the transcriptional cluster, but 3D phenotypes are buffered or redirected by compensatory increases in p53 signaling. Disabling p53 function synergizes with NRF2 deficiency, suppressing normal 3D proliferation and promoting irregular hyperproliferation in a transformed-yet-premalignant derivative. These observations are consistent with an integrated-systems model of NRF2–p53 signaling that encodes a shared oxidative-stress trigger and common pool of antioxidant target genes without any further crosstalk. Among clinical specimens, NRF2–p53 coordination is retained in normal primary breast tissue and hormone-negative DCIS. However, the two pathways are largely uncoupled in triple-negative breast cancers (TNBCs), where p53 is usually mutated (*35*). The integrated NRF2–p53 model predicts variable extents of uncoupling among TNBCs lines, and high uncoupling coincides with the most-severe 3D growth alterations upon NRF2 knockdown. Past work on NRF2 in breast cancer has focused on its direct interactions with TNBC tumor suppressors (*36, 37*). Our results suggest a broader systems-level role for NRF2 and p53 in oxidative-stress tolerance of normal breast–mammary epithelia and hormone-negative premalignancies.

## Results

### Statistical bioinformatics links gene-cluster regulation to NRF2 and p53

We began by looking within the gene cluster (Fig. 1A) for potential regulatory mechanisms. The only transcription factor in the cluster is *JUND*, and we showed previously that its chronic knockdown in MCF10A-5E cells (*20*) causes specific morphometric defects during spheroid growth (*24*). We revisited these results by acutely knocking down *JUND* with inducible shRNA and measuring transcript abundance of cluster genes by quantitative PCR (see Materials and Methods). Surprisingly, other than *JUND* itself, no transcripts were reliably altered by knockdown (fig. S1A), supporting a regulatory role for other factors outside of the cluster.

We constrained the search for candidate regulators by using maximum-likelihood inference (*38*) to estimate a frequency of bimodal transcriptional regulation (*39*) for the gene cluster. Given the 10-cell-averaged fluctuations from the original study (Fig. 1A) (*20*), the maximum-likelihood approach inferred two lognormal regulatory states defined by transcript abundance (Fig. 1B). The data supported a low-abundance regulatory state predominating in 58% of ECM-attached cells along with a second, high-abundance subpopulation in the remaining 42%. The frequency estimates placed quantitative bounds on the bimodal characteristics of upstream regulatory mechanisms.

Next, we applied a panel of bioinformatics approaches to search for transcription factors that might impinge upon the gene cluster (see Materials and Methods). The informatic methods adopt different strategies for assessing binding-site overrepresentation (*40–43*). Therefore, we intersected their respective outputs to arrive at predictions that were robust to algorithmic details. The analysis converged upon two transcription factors: the G1/S regulator E2F1 and the stress-response effector NRF2 (Fig. 1C). We assessed the relative activation of the NRF2 and E2F1 pathways in single cells by quantitative immunofluorescence for total stabilized NRF2 protein or hyperphosphorylated RB (pRB = disinhibited E2F1; see Materials and Methods). In 3D spheroid cultures, pRB immunostaining was bimodal, but high-pRB cells were far rarer than the inferred regulatory frequency of the gene cluster (Fig. 1D, upper). In 2D cultures, pRB staining was over twice as immunoreactive and nearly twice as prevalent in the population (Fig. 1D, middle). The reduced proportion of high-pRB cells in 3D is consistent with the proliferative suppression of late-stage spheroid cultures (*23*). A 3D-like distribution of pRB was achieved in 2D cultures upon addition of dilute ECM (Fig. 1D, lower) stemming from soluble proliferation-suppressing factors in the reconstituted basement membrane preparation (*44*). By contrast, NRF2 stabilization was only distinctly bimodal in 3D spheroids, and the observed frequency of low- and high-NRF2 states almost exactly coincided with that inferred for the gene cluster (Fig. 1E). The results built a strong statistical argument for NRF2 as a covarying regulator of the gene cluster.

The NRF2-associated gene cluster (Fig. 1A) was originally identified by quantitative analysis of transcriptomic fluctuations among 4557 genes profiled by oligonucleotide microarray (*20*). Recently, the same samples were reprofiled by 10-cell RNA sequencing (10cRNA-seq) (*45*), creating an opportunity to look more deeply at covariates with the NRF2-associated gene cluster. We used the median ranked fluctuations of the cluster across 10-cell samples (Fig. 1A) and surveyed the 10cRNA-seq data for genes that covaried (Spearman *ρ* > 0.5, *q* < 0.10), identifying 633 candidates (Fig. 2A). When this expanded cluster was assessed for functional enrichments by Gene Ontology (GO) (file S1) (*46*), we noted multiple GO terms linked to cell stress (“Response to stress”, “Oxidative stress”) and the transcription factor p53 (“DNA damage response”, “p53 pathway”; *q* < 0.05 by hypergeometric test). p53 is sporadically stabilized in regenerating epithelia such as the intestine and skin, but p53 activation in quiescent tissues is rare (*47*). Recognizing the residual proliferation observed in 3D cultures (Fig. 1D), we immunostained for p53 and found nonuniform stabilization associated with the abundance of NRF2 in single cells (Fig. 2B, estimated mutual information: MI = 0.15 [0.12–0.18]; see Materials and Methods). The analysis raised the possibility of a coordinated NRF2–p53 regulatory event triggered heterogeneously when breast epithelial cells proliferate and organize in reconstituted ECM.

**Fig. 2.**
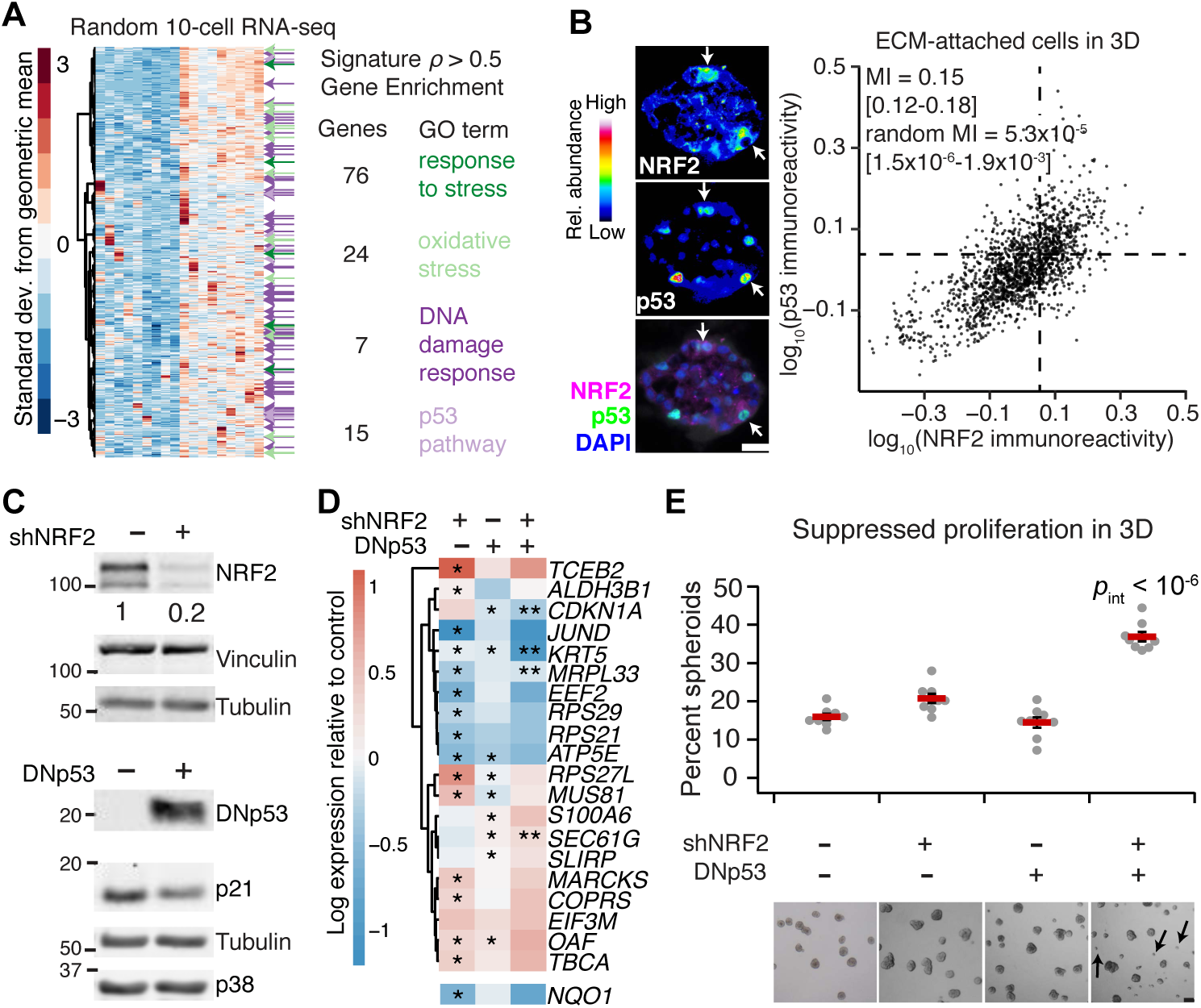
Transcriptome-wide covariate analysis of the NRF2-associated gene cluster suggests a coordinated adaptive-stress response involving p53. (**A**) Transcripts covarying with the median NRF2-associated fluctuation signature (Fig. 1A) measured by 10-cell RNA sequencing (*45*) of ECM-attached MCF10A-5E cells grown as 3D spheroids (*n* = 18 10-cell pools from GSE120261). Selected Gene Ontology enrichment analysis (green and purple) is shown for the transcripts with a Spearman correlation (*ρ*) greater than 0.5. The complete list of enrichments is available in file S1. (**B**) Stabilization of NRF2 and p53 proteins is coordinated in ECM-attached MCF10A-5E cells grown as 3D spheroids. Representative pseudocolored images for NRF2 (upper left) and p53 (middle left) are shown merged with DAPI nuclear counterstain (lower left). White arrows indicate concurrent NRF2 and p53 stabilization. Median-scaled two-color average fluorescence intensities are quantified (right) along with the log-scaled and background-subtracted mutual information (MI) with 90% CI for *n* = 1691 cells segmented from 50–100 spheroids from two separate 3D cultures. (**C**) Genetic perturbation of NRF2 by inducible shRNA knockdown (upper) and p53 by inducible expression of a FLAG-tagged carboxy terminal (residues 1-13, 302-390) dominant-negative p53 (DNp53, lower) (*49*). MCF10A-5E cells were treated with 1 µg/ml doxycycline for 72 hr (upper) or 24 hr (lower) and immunoblotted for NRF2 or FLAG with vinculin, tubulin, and p38 used as loading controls and p21 used to confirm efficacy of DNp53. The negative control for shNRF2 was an inducible shGFP, and the negative control for DNp53 was FLAG-tagged LacZ. (**D**) Single and combined perturbations of NRF2 and p53 have complex effects on the associated gene cluster (Fig. 1A). *NQO1* was used as a control for efficacy of shNRF2, *CDKN1A* shows efficacy of DNp53. MCF10A-5E cells with or without NRF2 knockdown or DNp53 were treated with 1 µg/ml doxycycline for 48 hr, grown as 3D spheroids for 10 days, and profiled for the indicated genes by quantitative PCR. Data are shown as the log_2_ geometric mean relative to the negative control (shGFP + FLAG-tagged LacZ), with asterisks indicating significant changes (left and middle columns) or interaction effects (right column) by two-way ANOVA of *n* = 8 independent 3D-cultured samples and a false-discovery rate of 5%. The complete set of transcripts in the gene cluster is shown in fig. S1C. (**E**) Dual inactivation of NRF2 and p53 causes synergistic proliferative suppression in MCF10A-5E 3D spheroids. Black arrows indicate proliferation-suppressed spheroids. Data are shown as the mean percentage of proliferation-suppressed spheroids ± s.e.m. of *n* = 8 independent 3D-cultured samples after 10 days. Statistical interaction between NRF2 and p53 (*p_int_*) was assessed by two-way ANOVA with replication. Scale bars are 20 µm (B) and 100 µm (E).

NRF2 co-immunoprecipitates with p53 in triple-negative breast cancer cells harboring gain-of-function p53 mutations, but this complex is absent in MCF10A cells with wildtype p53 (*37*). Loss of wildtype p53 function in MCF10A cells yields only minor 3D culture defects, but gain-of-function p53 mutants strongly perturb 3D architecture (*48*). Suspecting that some of p53’s effects could be explained through NRF2, we inducibly knocked down NRF2 with shRNA and inducibly coexpressed a truncated p53 (*49*) that acts as a dominant negative (DNp53; Fig. 2C). Compared with the gene-cluster response to JUND knockdown or constitutive E2F1 activation through RB inhibition with overexpressed human papillomavirus E7 protein, we observed substantially more alterations upon NRF2 knockdown (66%) or inhibition of p53 (31%; Fig. 2D and fig. S1, B to D). Compound perturbation of NRF2 and p53 elicited further nonadditive changes to multiple genes in the cluster, including synergistic reduction of the cyclin-dependent kinase inhibitor, *CDKN1A*, and the basal cytokeratin, *KRT5* (interaction *p* < 0.01 by two-way ANOVA). Although p53 can antagonize certain NRF2 target genes in reporter assays (*50*), significant antagonism was detected for only one transcript in the cluster (*MRPL33*, fig. S1C). Phenotypically, disruption of NRF2 reduced mean 3D growth by 10–13% (fig. S2), but dual perturbation with p53 gave rise to a surprising increase in aborted spheroids unable to grow in the culture (Fig. 2E). The penetrance of the phenotype (37%; range: 34–44%) was remarkably close to the percentage of cells showing stabilized NRF2 at the same time point of 3D culture (43%, Fig. 1E). For this clonal basal-like breast epithelial line (*20*), we conclude that 3D culture heterogeneously elicits NRF2- and p53-inducing stresses, which must be withstood for extended proliferation.

### NRF2 disruption in basal-like premalignancy causes similar p53 adaptations but different 3D phenotypes

We next asked how the cellular, molecular, and phenotypic relationships between NRF2 and p53 change in basal-like premalignancy by using isogenic MCF10DCIS.com cells (*51*) as a proxy for ductal carcinoma in situ (*52*). MCF10DCIS.com cells express oncogenic HRAS (*53*) and hyperproliferate as 3D spheroids (fig. S3A), but they retain wildtype p53 function, albeit at reduced levels compared to parental MCF10A cells (fig. S3, B and C). By two-color immunostaining, we found that NRF2–p53 co-stabilization was even more pronounced in MCF10DCIS.com cells (Fig. 3A, MI = 0.30 [0.27–0.33]). To identify common adaptive programs downstream of NRF2 deficiency, we inducibly knocked down NRF2 and profiled 3D spheroids by RNA sequencing (see Materials and Methods). Among transcripts consistently increased or decreased in both MCF10A-5E and MCF10DCIS.com spheroids, there was a significant enrichment in gene signatures encompassing p53, including transcriptional programs downstream of BRCA1, ATM, and CHEK2 (Fig. 3B and file S2). Consistent with these results, NRF2 knockdown in MCF10DCIS.com cells was sufficient to stabilize p53 significantly (fig. S4A). Stabilization of wildtype p53 upon NRF2 knockdown was also observed in premalignant *CHEK2^1100delC^* SUM102PT cells (*54*) and became even more pronounced when these cells were reconstituted with inducible wildtype CHEK2 (fig. S4B,C) (*55*). Thus, NRF2 impairment promotes p53 pathway activity in basal-like breast epithelia without the need for specific oncogenic drivers.

**Fig. 3.**
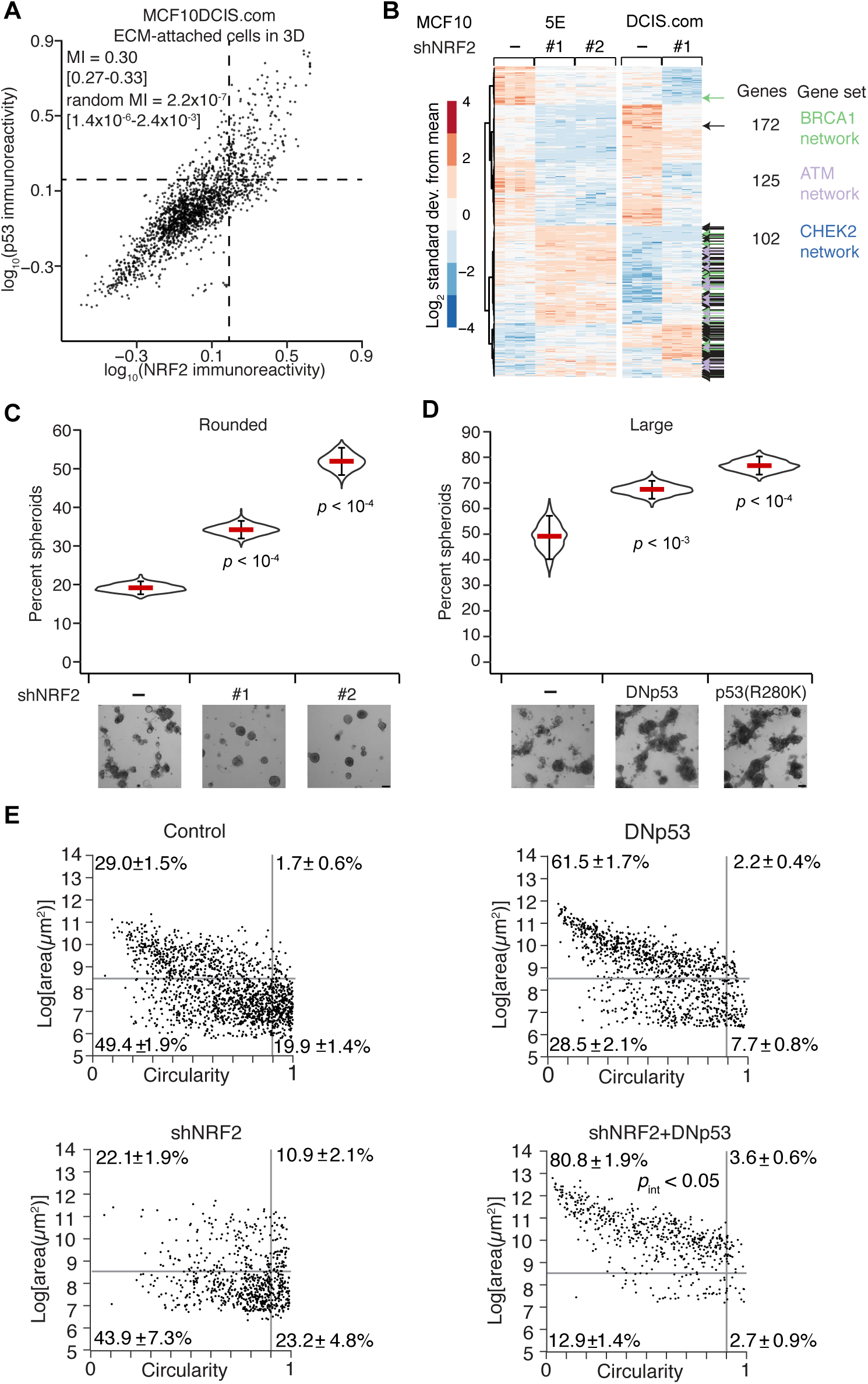
NRF2–p53 co-stabilization is enhanced and shNRF2-induced p53 adaptations are preserved in basal-like premalignancy but have different morphometric consequences. (**A**) Stabilization of NRF2 and p53 proteins is coordinated in ECM-attached MCF10DCIS.com cells grown as 3D spheroids. Median-scaled two-color average fluorescence intensities are quantified along with the log-scaled and background-subtracted mutual information (MI) with 90% CI for *n* = 1832 cells segmented from 70–110 spheroids from two separate 3D cultures. (**B**) Common changes in transcript abundance identified by RNA sequencing of MCF10A-5E (5E) and MCF10DCIS.com (DCIS.com) cells grown as 3D spheroids with or without NRF2 knockdown. The negative control for shNRF2 was an inducible shGFP (5E) or shLacZ (DCIS.com). Data are shown as log_2_-transformed Z-scores for genes detected at >5 transcripts per million from *n* = 4 biological replicates. Enriched gene sets for the BRCA1, ATM, and CHEK2 networks are indicated, with black denoting multiple enrichments. The complete list of enrichments is available in file S2. (**C**) NRF2 knockdown elicits a rounding phenotype in 3D-cultured MCF10DCIS.com cells. (**D**) p53 perturbation causes hyper-enlargement of 3D-cultured MCF10DCIS.com cells. (**E**) Dual inactivation of NRF2 and p53 synergistically increases the percentage of non-spherical, hyper-enlarged structures in 3D-cultured MCF10DCIS.com cells. For (C) to (E), cells with or without inducible perturbations were treated with 1 µg/ml doxycycline for 48 hr, grown as 3D spheroids for 10 days, imaged by brightfield microscopy, and segmented. For (C) and (D), data are shown as the mean ± 90% bootstrap-estimated CI from *n* = 8 biological replicates. For (E), data are shown as the mean ± s.e.m. of *n* = 8 biological replicates. Statistical interaction between NRF2 and p53 perturbations (*p_int_*) was assessed by two-way ANOVA with replication. Scale bars are 100 µm.

Despite many transcriptomic alterations in common with MCF10A-5E cells (Fig. 3B), MCF10DCIS.com cells yielded very different 3D phenotypes when NRF2 or p53 were perturbed. NRF2 knockdown did not detectably alter 3D growth (fig. S5A) but instead gave rise to more round, organized MCF10DCIS.com spheroids of high circularity compared to control (Fig. 3C), which reverted upon addback of an RNAi-resistant NRF2 mutant (fig. S5B). NRF2 deficiency also increased rounding in 3D cultures of SUM102PT cells with or without CHEK2 reconstitution (fig. S5C). By contrast, p53 disruption in MCF10DCIS.com cells with either DNp53 or a gain-of-function p53^R280K^ mutant increased the prevalence of hyperenlarged outgrowths (Fig. 3D). Combined NRF2–p53 perturbation elicited a synergistic increase in non-spherical hyper-enlargement (Fig. 3E, interaction *p* < 0.05 by two-way ANOVA), starkly contrasting the proliferative suppression observed with the same combination in nontransformed MCF10A-5E cells (Fig. 2E). The data suggested that the coordinate transcriptional adaptations of NRF2 and p53 are conserved in premalignant cells but insufficient to buffer the cellular phenotypes caused by single-gene perturbations in either pathway.

### NRF2 and p53 are coordinately stabilized by sporadic oxidative stress

Coordination of the NRF2–p53 pathways could be achieved if they shared the same inducer. We thus considered various potential upstream-and-intermediate triggers for NRF2 and p53 stabilization in basal-like breast epithelia. Inhibition of KEAP1 with the electrophile sulforaphane (*56*) stabilized NRF2 but not p53, and pharmacologic inhibition of MDM2 with nutlin-3 (*57*) stabilized p53 but not NRF2 (fig. S3, B to D), suggesting they act as parallel pathways downstream of a common inducer. An obvious candidate was DNA damage given *CDKN1A* and *MUS81* in the gene cluster (Fig. 1A) and the most-recognized function of p53 (*58*). However, chemotherapy-induced double-strand breaks did not appreciably stabilize NRF2 in cells with wildtype p53 (Fig. 4A), and genetically driving hyperproliferation (*59*) did not detectably impact regulation of the gene cluster in 3D spheroids (fig. S1, B and D). The lack of NRF2–p53 co-induction by conventional agonists prompted a search for less canonical activators.

**Fig. 4.**
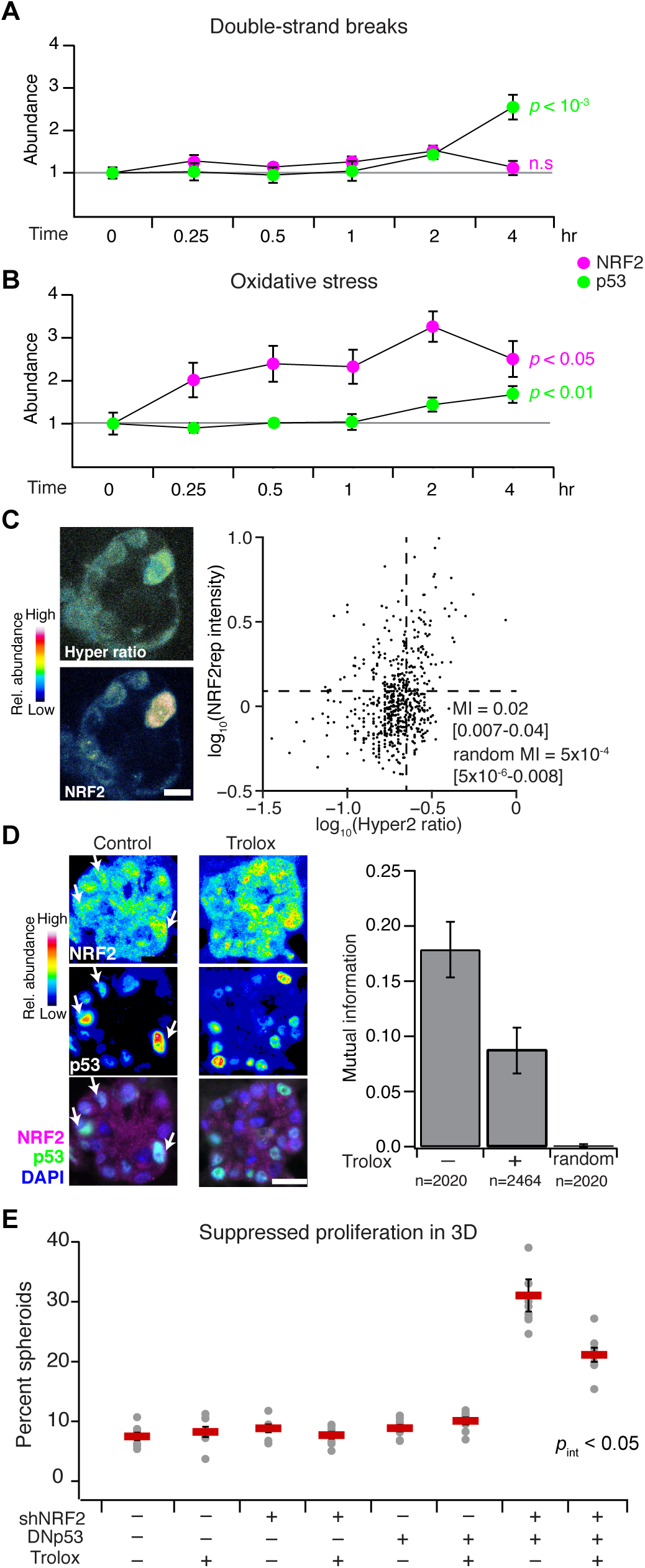
NRF2–p53 signaling coordination and 3D phenotypes arise from spontaneous and oncogene-induced oxidative stress. (**A** and **B**) NRF2 and p53 are jointly activated by oxidative stress but not by DNA double-strand breaks. MCF10A-5E cells were treated with 5 µM doxorubicin (double-strand breaks) or 200 µM H_2_O_2_ (oxidative stress) for the indicated time points, and NRF2 (magenta) or p53 (green) protein abundance was estimated by quantitative immunoblotting. Data are shown as the mean ± s.e.m. of *n* = 3 (A) or 4 (B) independent perturbations. (**C**) Endogenous oxidative stress is associated with NRF2 stabilization in 3D spheroids. MCF10A-5E cells stably expressing HyPer-2 (*65*) and mRFP1-NRF2 reporter (NRF2rep) were grown as 3D spheroids for 10 days and imaged by laser-scanning confocal microscopy. Representative pseudocolored images for HyPer-2 ratio (upper left) and mRFP1-NRF2 reporter (lower left) are shown. HyPer-2 ratios and mRFP1-NRF2 reporter fluorescence are quantified (right) along with the log-scaled mutual information (MI) with 90% CI for *n* = 605 cells segmented from 10–25 spheroids from four separate 3D cultures. (**D**) The antioxidant Trolox suppresses endogenous NRF2–p53 coordination during 3D culture. Representative pseudocolored images for NRF2 (upper left) and p53 (middle left) are shown merged with DAPI nuclear counterstain (lower left). White arrows indicate concurrent NRF2 and p53 stabilization. The log-scaled and background-subtracted MI (right) is shown with 90% CI estimated from *n* = 1000 bootstrap replicates. (**E**) Trolox interferes with the synergistic proliferative suppression caused by dual inactivation of NRF2 and p53 in MCF10A-5E cells. Data are shown as the mean percentage of proliferation-suppressed spheroids ± s.e.m. of *n* = 8 independent 3D-cultured samples after 10 days. The overall effect of Trolox on spheroid size is shown in fig. S8. Statistical interaction between Trolox and NRF2–p53 (*p_int_*) was assessed by three-way ANOVA with replication. For (D) and (E), MCF10A-5E cells cultured for 10 days in 3D with or without 50 µM Trolox supplemented every two days. Scale bars are 10 µm (C) and 20 µm (D).

One shared inducer of the KEAP1–NRF2 and ATM–CHEK2–p53 pathways is oxidative stress (*60, 61*). In human breast tissue, elevated levels of reactive oxygen species are generated and tolerated by basoluminal progenitors (*62*), which are the cells of origin for basal-like breast cancer (*63*). We documented local niches of Nrf2 stabilization in the murine mammary gland during puberty (fig. S6), potentially linking NRF2 and oxidative stress in expanding progenitor(-like) cells, such as MCF10A. When MCF10A-5E cells were exogenously stimulated with H_2_O_2_, NRF2 was rapidly stabilized and, importantly, p53 also accumulated after several hours (Fig. 4B). Recognizing oxidative-stress heterogeneities in 3D spheroids (*21, 22, 64*), we used the genetically-encoded sensor HyPer-2 (*65*) together with a novel mRFP1-NRF2 reporter (NRF2rep) to colocalize intracellular H_2_O_2_ with stabilized NRF2 (see Materials and Methods and fig. S7). We observed a small-but-nonzero mutual information between HyPer-2 fluorescence ratios and NRF2rep (Fig. 4C, MI = 0.05 [0.02–0.10]; randomized MI = 0.0004 [0.0001–0.0007]), suggesting a weak (or complex) connection between the two. Next, we evaluated whether oxidative stress resided upstream of NRF2–p53 coordination by using the vitamin E analog Trolox to quench reactive oxygen species in the 3D cultures. Trolox treatment halved the mutual information between stabilized NRF2–p53 and significantly reduced the synergistic proliferative suppression caused by dual perturbation of NRF2 and p53 (Fig. 4, D and E, and fig. S8) Together, the data strongly suggested that NRF2 and p53 pathway co-regulation involves upstream heterogeneities in oxidative stress.

### An integrated NRF2–p53 model of oxidative stress reconciles pathway coordination with 3D phenotypes

To connect NRF2 and p53 co-stabilization with spontaneous heterogeneities in oxidative stress, we assembled an integrated computational-systems model. The model expands or condenses isolated modules of NRF2 and p53 signaling from the literature, fusing them through known or reported mechanisms of crosstalk and convergence (Fig. 5A). For the NRF2 pathway, we streamlined the detailed model of Khalil *et al.* (*66*) at several points. Instead of relying on ill-defined kinetic parameters for KEAP1-mediated ubiquitination, KEAP1:NRF2 complexes were modeled as separate oxidized or reduced species with distinct half-lives estimated by experiment (see Materials and Methods). We likewise abandoned the elaborate multistep encoding of thioredoxins, peroxiredoxins, and glutathione transferases (*66*) by substituting a simpler, lumped pool of antioxidant enzymes in the model. The resulting architecture is similar to the general negative-feedback control scheme of stress-response gene-regulatory networks described by Zhang & Anderson (*67*). Last, we retained the nucleocytoplasmic trafficking of stabilized NRF2 to account for observations that H_2_O_2_ stimulation retains NRF2 in the cytoplasm longer than treatment with the electrophilic stress, sulforaphane (fig. S9). Oxidative stress feeds directly into the NRF2 module according to a basal production rate of reactive oxygen species (ROS), which was adjusted in the final model to yield steady-state intracellular H_2_O_2_ concentrations consistent with the literature (*68*).

**Fig. 5.**
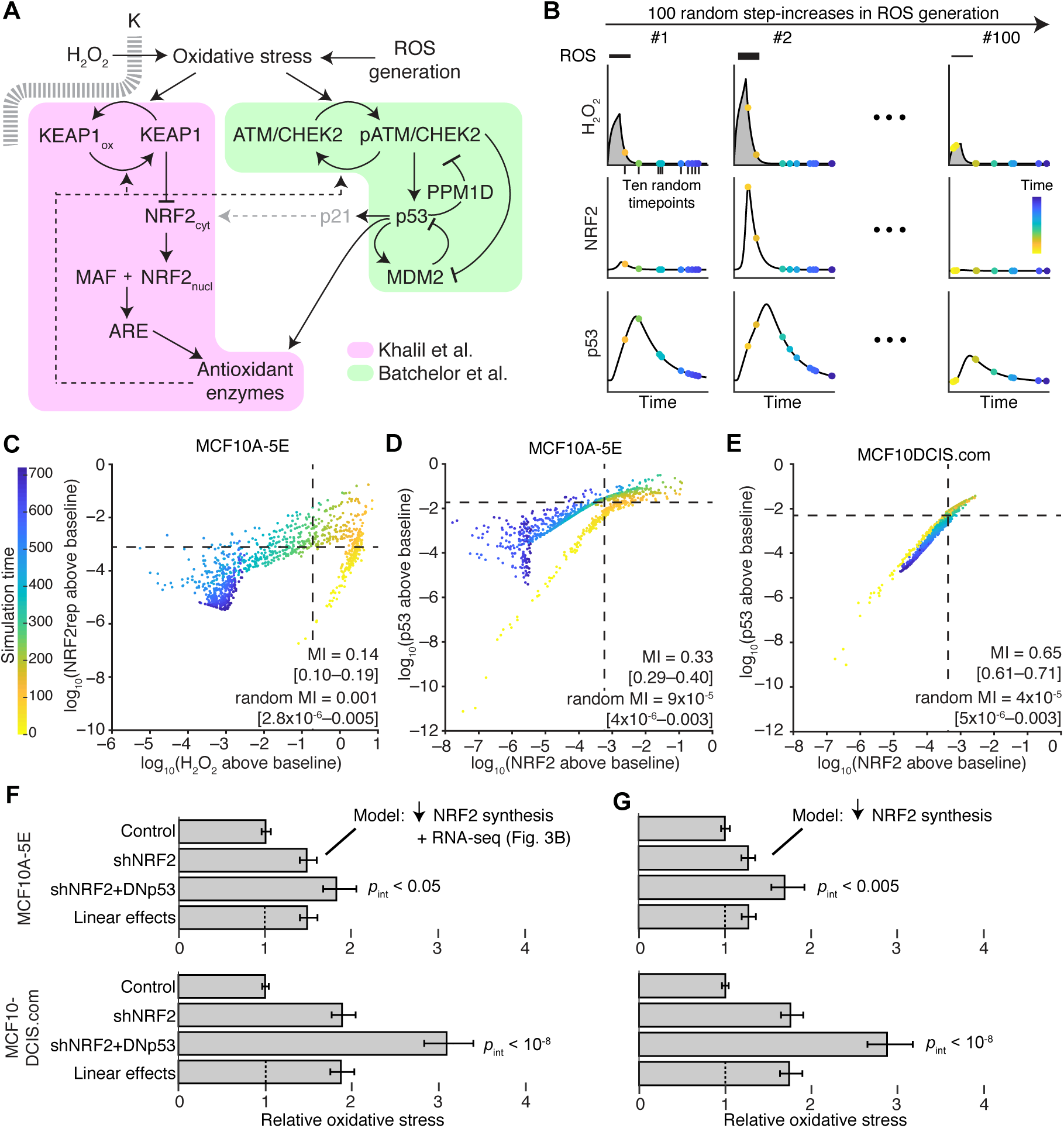
NRF2–p53 pathway coordination and synergistic phenotypes are captured by an integrated-systems model of oxidative stress. (**A**) Connecting NRF2 and p53 signaling models (*66, 67, 69*) through oxidative-stress activators and antioxidant target enzymes. Additional crosstalk linking p53 to NRF2 through p21 (*71*) is conditionally incorporated (gray). (**B**) Simulation strategy for quantifying association between signaling intermediates. The model was challenged with various ROS production rates and randomly sampled at multiple intermediate time points (yellow to blue). Integrated intracellular H_2_O_2_ (gray) is used for phenotype predictions related to NRF2 and p53 perturbation. (**C**) Intracellular H_2_O_2_ concentration is associated with a reporter of NRF2 stabilization (NRF2rep). (**D** and **E**) Coordination of NRF2 and p53 stabilization is high in the oxidative-stress model and increases further in simulations of premalignancy. (**F** and **G)** Modeling NRF2 knockdown by reduced synthesis captures the synergistic oxidative-stress profile of cells harboring dual perturbation of the NRF2 and p53 pathways. In (F), transcriptional changes secondary to NRF2 knockdown were added to the model according to the results in Fig. 3B. For (C) to (E), simulated time points are shown as the log-scaled and background-subtracted mutual information (MI) with 90% CI for ten time points from *n* = 100 random ROS generation rates. For (F) and (G), time-integrated intracellular H_2_O_2_profiles are scaled to the unperturbed simulations and shown as the mean oxidative stress with 90% CI from *n* = 100 random ROS generation rates.

For the p53 pathway, we built upon the base model of Batchelor *et al.* (*69*), which was originally used to describe oscillations in p53 abundance after ionizing radiation. In this model, the kinases ATM and CHEK2 act as aggregate sensors–transducers of the DNA damage response (Fig. 5A). They phosphorylate and stabilize p53 against degradation triggered by the ubiquitin ligase MDM2, which is also directly phosphorylated and inactivated by ATM. Stabilized p53 promotes its own degradation by inducing *MDM2* transcripts and deactivates ATM–CHEK2 by enhancing transcription of the phosphatase *PPM1D*. For the integrated model, oxidative stress replaced DNA double-strand breaks as the pathway trigger, recognizing that ATM autoactivates in the presence of oxidants (*61*). Further, in response to oxidative stress, proper induction of many antioxidant enzymes requires p53 (*70*), which contributes to the overall antioxidant pool along with ARE target genes (Fig. 5A). As a final candidate for NRF2–p53 crosstalk, we considered reports that the p53 target gene, p21 (CDKN1A), directly stabilizes NRF2 by interfering with KEAP1-catalyzed turnover (*71, 72*). Together, the modifications provided an integrated model of NRF2–p53 signaling downstream of oxidative stress with enough molecular detail to enable kinetic and functional predictions.

We revisited the oxidative stress time course (Fig. 4B) to append immunoblot quantification of ATM–CHEK2 phosphorylation and p21 abundance after H_2_O_2_ addition (fig. S10A). Exogenous H_2_O_2_ was encoded as an extracellular spike-in that decayed rapidly and spontaneously (*73*) amidst a basal ROS generation rate yielding a realistic intracellular H_2_O_2_ burden at steady state (*68*). The H_2_O_2_ partition coefficient in the model was calibrated to capture the magnitude of NRF2 stabilization (see Materials and Methods). Likewise, the parameters for H_2_O_2_-induced autoactivation of ATM–CHEK2 and signal inactivation were defined to align with the time-delayed kinetics and duration of p53 stabilization (fig. S10B). In this model, addition of p53–p21–NRF2 crosstalk (*71*) caused NRF2 stabilization to peak earlier and deactivate faster than observations (fig. S10C). We were also unable to detect even transient short-range interactions between inducible BirA*-fused versions of p21 or NRF2 and endogenous NRF2 or p21 by proximity ligation (fig. S11). The results thus argued against p53–p21–NRF2 crosstalk during oxidative stress in these cells.

With the provisionally calibrated base model, we sought to test whether the encoded mechanisms of regulation were sufficient to capture prior observations relating NRF2, p53, and oxidative stress. The data obtained by quantitative fluorescence microscopy (Fig. 2B, 3A, and 4C) presumably arose from spontaneous oxidative stress that was occurring transiently and asynchronously during imaging. We mimicked oxidative-stress transients by triggering a step increase in the rate of ROS production for two hours followed by relaxation of the system for an additional 10 hours (Fig. 5B). The magnitude of the step was sampled lognormally to elicit intracellular H_2_O_2_ concentrations within the range of HyPer-2 ratios observed experimentally (see Materials and Methods). We represented the asynchrony of image acquisition by randomly selecting ten snapshots of the network for each model iteration. This collection of 1000 snapshots (100 random generation rates x 10 random time points) was used to quantify coordination of species within the model.

For connecting oxidative stress to NRF2 stabilization (Fig. 4C), we expanded the base model to include the mRFP1-NRF2 reporter, which does not bind DNA or interact with MAF proteins and requires ~1 hr to mature fully (*74*) (see Materials and Methods). By contrast, HyPer-2 becomes fully oxidized within ~1 min of H_2_O_2_ addition (*65*), enabling intracellular H_2_O_2_ concentration in the model to be used directly as a surrogate of HyPer-2 fluorescence ratio. We calculated the mutual information from 1000 simulated snapshots and found that the two reporters were statistically coupled in the model (Fig. 5C, MI = 0.14 [0.10–0.19]; randomized MI = 0.001 [2.8×10^−6^–0.0005]). Associations were stronger than those observed by experiment (Fig. 4C) due to the early time points sampled in the model (yellow-orange times in Fig. 5C), suggesting that peak H_2_O_2_ transients may be difficult to observe in practice. We next compared endogenous NRF2–p53 co-stabilization between MCF10A-5E and MCF10DCIS.com cells. The base MCF10A-5E model was adjusted to reflect i) proportional differences in species abundance estimated from RNA-seq (see Materials and Methods) and ii) an increased ROS generation rate estimated from HyPer-2 imaging (fig. S7D). NRF2–p53 mutual information was much less dependent on signaling transients, and coupling was substantially higher in MCF10DCIS.com cells. The simulations are consistent with immunofluorescence data (Fig. 3A, 4C, 5, D and E) and support that NRF2–p53 pathway kinetics are accurately encoded in the base model.

We asked whether the base model could also relate to the synergistic phenotypes observed upon dual NRF2–p53 perturbation in MCF10A-5E and MCF10DCIS.com cells (Fig. 2E, 3E). We mimicked shNRF2 by reducing the NRF2 production rate fivefold in the model (Fig. 2C) and encoding secondary transcriptional adaptations in other components by using the associated RNA-seq data (Fig. 3B). For DNp53, the p53 species was rendered unable to induce transcription of MDM2, PPM1D, p21, and its share of the antioxidant enzyme pool. After re-establishing steady state, the perturbed models were challenged with the random step increase in ROS production described above. We used the time-integrated intracellular H_2_O_2_ concentration as the overall measure of oxidative stress experienced during simulation with either the MCF10A-5E or MCF10DCIS.com initial conditions. For both cell lines, the base model predicted synergistic increases in oxidative stress beyond the linear superposition of shNRF2 and DNp53 effects (Fig. 5F). Encouragingly, the same conclusions were reached with models that simply encoded the reduced NRF2 production rate without secondary adaptations (Fig. 5G). Beyond oxidative-stress inducers and antioxidant target enzymes, we conclude that the NRF2– p53 network does not require any additional mechanisms to capture signaling coordination or phenotypic interactions.

### NRF2–p53 co-regulation occurs in normal breast tissue and hormone-negative DCIS but not invasive TNBC

The regulatory heterogeneities observed in 3D culture often reflect adaptations in hormone-negative premalignancy (*24*) that become further disrupted in TNBCs (*25*). We thus sought to quantify NRF2–p53 coordination in TNBC and premalignant DCIS lesions, using adjacent-normal tissue as a comparator. The *TP53* gene is frequently mutated in TNBC (*35*) and gives rise to loss of p53 protein or hyperstabilization of a dominant-negative mutant in tumors (*75*). By contrast, prior immunohistochemistry of NRF2 abundance in breast carcinomas was inconclusive (*76*), owing to an anti-NRF2 antibody that was later shown to be non-specific (*77*). There was an opportunity to revisit NRF2–p53 abundance heterogeneities from the perspective of co-stabilization, with a focus on TNBC and its precursor lesions.

Using a knockout-verified commercial antibody (*78*), we immunoblotted with our production lot and confirmed detection of basal and induced NRF2 with only ~35% immunoreactivity attributed to nonspecific bands (fig. S12, A to C). By immunohistochemistry, the antibody detected endogenous NRF2 stabilized with electrophiles in paraffin sections of cell pellets (fig. S12D). The antibody has also been used independently to track NRF2 abundance in other solid tumors (*79*). However, when we stained adjacent-normal epithelium immunohistochemically, NRF2 was not clearly discernible (Fig. 6A, upper). In MCF10A 3D spheroids, stabilized p53 is not detected by immunohistochemistry either (*80*), and yet we readily visualized it by immunofluorescence (Fig. 2B). Therefore, to improve signal-to-background and facilitate multiplex quantification, we used two-color immunofluorescence after antigen retrieval, segmenting 24,949 normal and transformed epithelial cells in 15 cases of TNBC and hormone-negative DCIS.

**Fig. 6.**
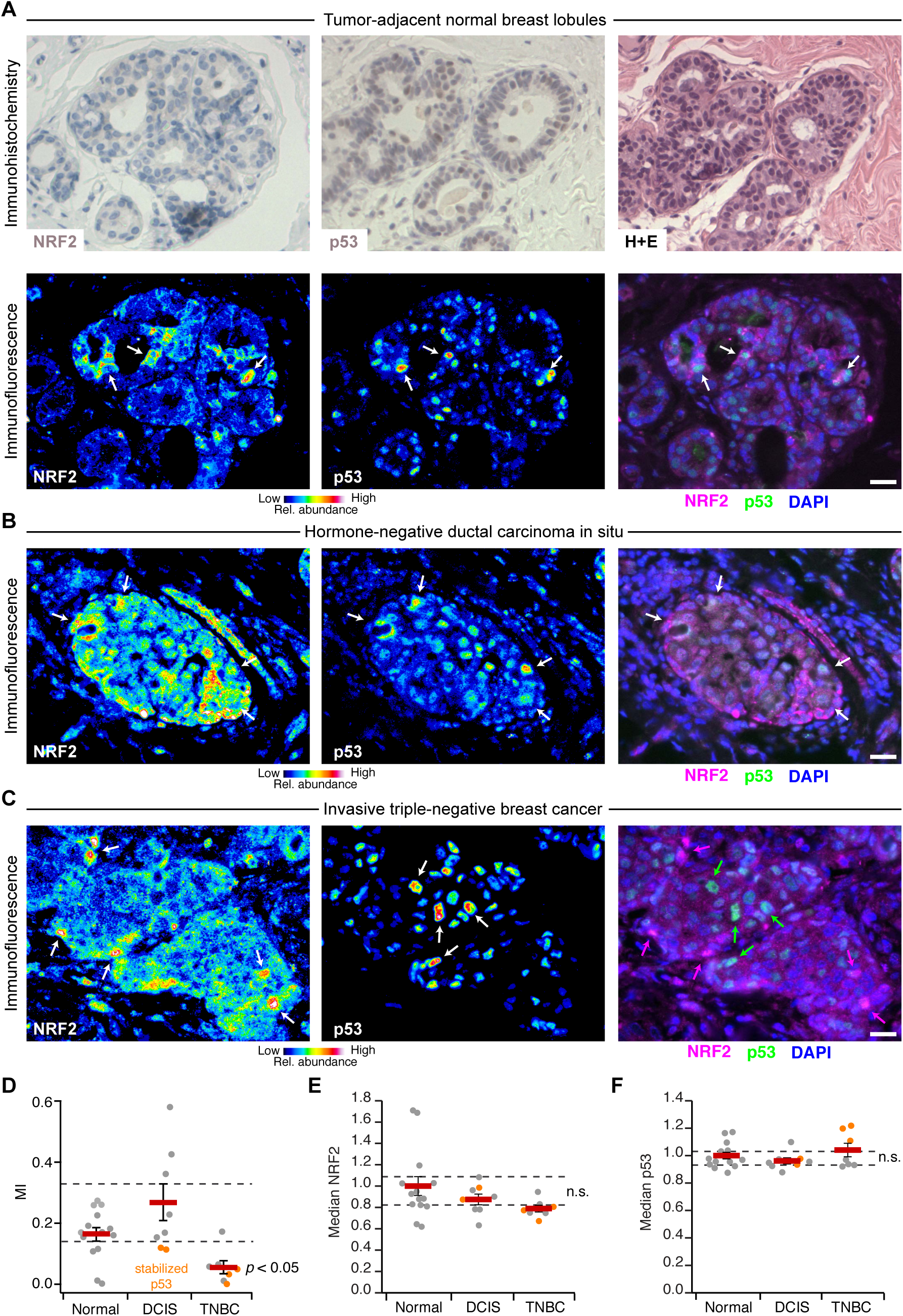
NRF2 and p53 are co-stabilized in breast epithelial tissue and premalignant lesions but uncoupled in triple-negative breast cancer. (**A**) Immunohistochemistry (upper) and immunofluorescence (lower) for NRF2 and p53 in tumor-adjacent normal breast lobules. Hematoxylin and eosin (H+E, upper right) histology is from a serial paraffin section for p53. Images from a tumor-adjacent normal breast duct are shown in fig. S13. (**B** and **C**) Multicolor immunofluorescence for NRF2 and p53 in (B) hormone-negative ductal carcinoma in situ and (C) triple-negative breast cancer. (**D**) NRF2–p53 mutual information (MI) is lost in triple-negative breast cancer. (**E** and **F**) Overall NRF2 and p53 immunoreactivity is not consistently altered during triple-negative breast cancer progression. n.s., not significant (*p* > 0.05). For (A) to (C), immunofluorescence is shown as representative pseudocolored images for NRF2 (left) and p53 (middle) are shown merged with DAPI nuclear counterstain (right). White arrows indicate concurrent NRF2 and p53 stabilization, and magenta or green arrows indicate stabilization of NRF2 or p53 separately. Scale bars are 20 µm. For (D) to (F) data are shown as the mean ± s.e.m. of *n* = 14 cases with tumor-adjacent normal epithelium (Normal), 8 cases with ductal carcinoma in situ (DCIS), and 7 cases of triple-negative breast cancer (TNBC).

In adjacent-normal epithelium, we observed local niches of stabilized NRF2 in lobules and ducts, which often corresponded with stabilized p53 (Fig. 6A, lower, and fig. S13). Stabilized NRF2 was frequently detected in the cytoplasm, consistent with the prolonged cytoplasmic localization observed in H_2_O_2_-treated cells compared to cells stressed with an electrophile (fig. S9). The results corroborated recent findings that KEAP1 senses oxidative stress differently than electrophilic stress (*60*). The patterns of NRF2–p53 co-accumulation were largely preserved in hormone-negative DCIS (Fig. 6B and fig. S14), even in cases with hyperstabilized p53 that was likely mutated (see below). Nuclear localization of NRF2 was also more prominent, perhaps reflecting the stronger ROS generation rates of transformed cells (*81*). Strikingly, NRF2 and p53 were almost completely uncoupled in invasive TNBCs (Fig. 6C and fig. S14), reflecting a profound shift in single-cell regulation. We quantified NRF2–p53 coordination by mutual information and found that it was largely eliminated in regions of invasive TNBC, irrespective of whether p53 was chronically stabilized or not (Fig. 6D). Such alterations were not apparent in regional estimates of protein abundance by cell population-averaged fluorescence, where neither NRF2 nor p53 were reproducibly different among groups (Fig. 6, E and F). We conclude that 3D culture in reconstituted basement membrane co-stimulates the NRF2–p53 pathways akin to that observed in normal breast tissue and hormone-negative premalignancy. Full-blown TNBC, by contrast, evokes a different set of dependencies.

### TNBC adaptations to p53 disruption predict variable NRF2 miscoordination, NRF2-deficient oxidative-stress profiles, and 3D growth responses

*TP53* is the most-frequently mutated gene in TNBC (*35*), and transcriptomic analyses support it as a prevalent founder mutation in the disease (*82*). Disrupting p53 would undoubtedly impact transcriptional feedback and the overall cellular response to oxidative stress (Fig. 5A). Conversely, neither *NFE2L2* nor *KEAP1* are mutated in breast cancer (*83*), but unclear is whether wildtype NRF2 might serve as a transient “non-oncogene” (*84*) that promotes stress tolerance during early tumorigenesis. Compared to in situ lesions, the stromal environment of invasive tumors is stiffer and more mesenchymal (*85*), which may render NRF2 signaling dispensable at later stages. We wondered whether the fragmentation of the NRF2–p53 network in TNBC cells and its origins could be reconciled with the systems model.

Using RNA-seq data from the NIH LINCS consortium (*86*) on 15 TNBC lines with mutated p53 (six claudin-low subtype, nine basal-like subtype), we adjusted initial conditions from the original MCF10A model and removed all transcriptional processes downstream of p53 (Fig. 7A; see Materials and Methods). The individual TNBC models were run to steady state and then challenged with increased ROS generation rates as in Fig. 5B. The coordination between NRF2 and mutant p53 was calculated by mutual information, and the integrated H_2_O_2_response was scaled to that of MCF10A-5E cells as a relative measure of ROS tolerance. The goal was to associate the model-derived predictions with NRF2-knockdown phenotype in ROS-generating environments such as 3D culture. To the extent possible, we hoped that 3D growth in reconstituted basement membrane might quantify any vestigial requirements for NRF2 signaling from the in situ stage of the TNBC lines.

**Fig. 7.**
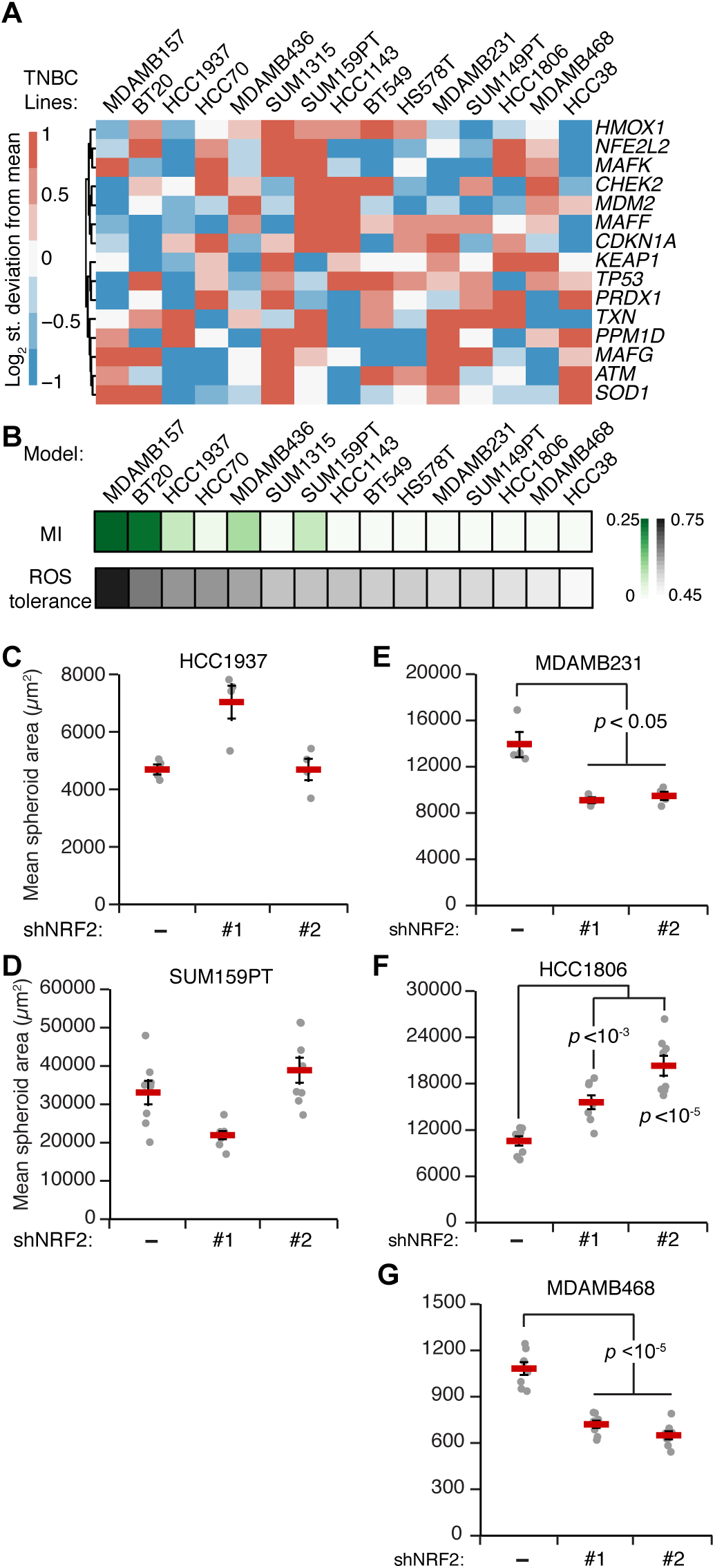
TNBC-specific signatures of the oxidative-stress network predict NRF2–p53 coupling and the response to NRF2 perturbations. (**A**) Transcripts per million for the indicated TNBC cell lines scaled to MCF10A cells from the NIH LINCS dataset (*86*). The clustered transcripts were used to adjust the initial conditions of the model simulations for each cell line. (**B**) NRF2–p53 mutual information (MI) correlates with ROS tolerance in TNBC simulations. ROS tolerance was defined as the integrated intracellular H_2_O_2_ concentration in each cell line compared to that of MCF10A-5E cells in response to an increased ROS production rate as in Fig. 5B. (**C** and **D**) 3D growth of TNBC cells with higher ROS tolerance and NRF2-p53 MI is unaffected by NRF2 knockdown. (**E** to **G**) TNBC cells with lower ROS tolerance and NRF2-p53 coordination show changes in 3D growth upon NRF2 knockdown. For (C) to (E), data are shown as the mean ± s.e.m. of *n* = 4–8 biological replicates.

For all TNBC lines, the model predicted significantly reduced covariation between mutant p53 and NRF2 compared to MCF10DCIS.com cells with wildtype p53 (Fig. 7B, MI < 0.25). We noted a spectrum of residual NRF2–p53 co-stabilization from weak (e.g., HCC1937, SUM159PT) to virtually nonexistent (MDA-MB-468, MDA-MB-231). Despite complete p53 deficiency in the model, this residual NRF2–p53 mutual information correlated strongly with the relative increase in oxidative stress upon simulated NRF2 knockdown (Fig. 7B). Neither of these predictions mapped directly to specific transcripts in the TNBC-specific RNA-seq data (Fig. 7A), reinforcing that the models were making nonobvious predictions about oxidative-stress handling.

To connect the model predictions with a continued role for NRF2 signaling in TNBC behavior, we selected five lines along the spectrum of mutual information and ROS tolerance. HCC1937 and SUM159PT cells were both predicted to have residual NRF2–p53 coordination and moderate ROS tolerance (Fig. 7B). Accordingly, inducible knockdown of NRF2 in these lines did not lead to any consistent changes in 3D growth (Fig. 7, C and D). By contrast, MDA-MB-231, HCC1806, and MDA-MB-468 cells were predicted to have among the least NRF2–p53 co-stabilization and ROS tolerance (Fig. 7B). Knockdown of NRF2 in these lines with two different shRNAs caused significant increases or decreases in overall cell growth (Fig. 7, E to G). Thus, model and experiment support that, despite p53 mutation, residual NRF2–p53 coupling indicates the primordial susceptibility of triple-negative malignancies to perturbations in the NRF2 pathway.

## Discussion

This work posits ROS as an endogenous, spatially heterogeneous trigger of dual NRF2–p53 activation in breast–mammary epithelia surrounded by basement membrane ECM. NRF2 and p53 regulate target-gene abundance—both cooperatively and independently—to promote stress tolerance and adaptation. NRF2 deficits are buffered by compensatory increases in p53 signaling, and dramatic ROS-dependent phenotypes arise when both pathways are perturbed. In hormone-negative premalignant lesions, stabilization of NRF2–p53 remains coordinated, even in cases where p53 has likely mutated. At this preinvasive stage, NRF2 should be most important for tumorigenesis. After invasion through basement membrane and progression to TNBC, the stromal microenvironment reduces overall NRF2 signaling and uncouples it from (now-mutant) p53. Here, the effect of activating or inhibiting NRF2 will depend more on the exact cellular context and thus be unpredictable (for example, Fig. 7, E to G). Despite the overall complexity of NRF2- and p53-mediated transcriptional programs (*87, 88*), the coordinated response to oxidative stress is captured by a relatively simple mathematical encoding. Known core mechanisms of NRF2–p53 regulation are brought together by a shared ROS inducer and a common pool of detoxifying target genes without the need for any further crosstalk. Therefore, oxidative-stress handling in normal breast–mammary epithelia is usefully abstracted as two stability-regulated transcription factors working independently toward a common homeostatic goal.

Although NRF2 is not an oncogene for breast cancer, it has been connected with multiple breast-cancer tumor suppressors previously. In mammary epithelial cells, loss of *Brca1* (a predisposing event for basal-subtype TNBC) destabilizes Nrf2 and causes an increase in ROS favoring the future acquisition of p53 mutations (*36, 89*). In human breast cancer cells, gain-of-function p53 mutants interact directly with NRF2 and may help retain NRF2 in the nucleus (*37*). If certain p53 mutations were also to promote NRF2 stabilization, then it would provide a two-for-one benefit to cancer progression by relieving tumor suppression and conferring ROS tolerance constitutively. However, we did not note any association between gain-of-function p53 mutants and NRF2 abundance in TNBC lines (fig. S12C), suggesting that KEAP1 regulation predominates as indicated by the TNBC models. Chronic activation of the NRF2 pathway (for example, by activating *NFE2L2* mutation or *KEAP1* loss) may be disfavored if elevated intracellular ROS is not permanent. The models suggest that supraphysiological activation of NRF2 would lead to runaway induction of antioxidant enzymes, causing reductive stress as documented for NRF2 in other tissues (*90*). Wildtype NRF2 function must be sufficient to buffer cells from the early stresses of premalignancy and p53 disruption, allowing invasive TNBCs to deactivate the pathway when it is no longer needed. There are parallels to FOXO transcription factors (*91*), which are reversibly inactivated by mitogenic signals yet provide critical oxidative-stress tolerance when the breast-cancer tumor suppressor RUNX1 is disrupted (*22, 92, 93*).

Breast cancer cell lines organize very differently in 3D culture (*94*), but their response to perturbations is often less disparate. For example, gain-of-function p53 mutations cause luminal filling in MCF10A 3D cultures (*48*), similar to the delay in mammary-gland involution observed with mutant p53 in vivo (*95*). Reciprocally, knockdown of mutant p53 in MDA-MB-468 cells promotes luminal hollowing (*96*). Among p53-mutant TNBC lines, the impact of NRF2 knockdown on 3D growth was nonuniform but explainable through the oxidative-stress profiles inferred from TNBC-specific systems models. The balance of complexity and tractability make 3D spheroid–organoid cultures a compelling platform for systems-level dissection of cell-state heterogeneity and early tumorigenesis.

The 3D behavior of breast–mammary cancer cells is highly dependent on the surrounding ECM (*97*). Invasive cancers no longer encounter basement membrane ECM but must have bypassed it upon progression to carcinoma. Interestingly, although multiple TNBC lines will grow as 3D colonies in reconstituted basement membrane, others cannot, suggesting a type of cellular “amnesia” toward that past encounter. For cancers that do grow in 3D, the use of reconstituted basement membrane (as a more normal-like microenvironment) may give rise to cellular changes reminiscent of premalignancy. We exploited these changes to evaluate the relative importance of NRF2 signaling in different TNBC backgrounds. There are likely other opportunities to examine hurdles of premalignancy by using basement-membrane 3D cultures. For 3D-organoid biobanks (*19*), however, it is a reminder that such cultures are not propagating the primary breast tumor but rather tumor-derived cells in a more-primitive state.

Cancer mutations engage and cooperate with cell signaling in ways that are not captured by DNA sequencing (*98*). The coupling of the NRF2 and p53 pathways described here provides a robust oxidative stress-handling network for glandular morphogenesis and maintenance. But, this same coupling creates a redundancy upon which p53 mutations can occur and neoplasms can evolve. Our results give pause to the nutraceutical use of sulforaphane as a potent NRF2 stabilizer (*99*)—in lung cancer, where *KEAP1–NRF2* mutations are common and *TP53* is secondary, antioxidants accelerate tumor progression (*100*). The extraordinary complexity of ROS generation and its cellular effects reinforce the value of modeling redox networks at a granularity suited to a given physiology or pathology (*101*).

## Materials and Methods

### Plasmids

shRNA targeting sequences from the RNAi consortium (*102*) were cloned into tet-pLKO.1-puro as previously described (*38*) for shLuc (TRCN0000072250, Addgene #136587), shNRF2 #1 (TRCN0000281950, Addgene #136584), shNRF2 #2 (TRCN0000284998, Addgene #136585), shJUND #1 (TRCN0000416347, Addgene #136581), shJUND #2 (TRCN0000416920, Addgene #136583).

For the mRFP1-NRF2 reporter (Addgene #136580), the DNA binding domain of NRF2 was mutated (C506S) along with four leucines (L4A) in the leucine zipper region of the bZIP domain by site-directed mutagenesis of the pBabe mRFP1-NRF2 hygro plasmid (Addgene #136579) originally prepared by subcloning into pBabe mRFP1 hygro. The RNAi-resistant (RR) version of NRF2 (Addgene #136522) was prepared by introducing four silent mutations into the sequence targeted by shNRF2 #1 in pEN_TT 3xFLAG-NRF2 (Addgene #136527). Site-directed mutagenesis was performed with the QuikChange II XL kit (Agilent).

pDONR223 CHEK2 was obtained from the human Orfeome V5.1 (*103*). CHEK2 amplicon was prepared with XbaI and MfeI restriction sites and cloned into pEN_TTmiRc2 3xFLAG (Addgene #83274) that had been digested with SpeI and MfeI (Addgene #136526). BirA* was cloned out of pcDNA3.1 mycBioID (Addgene) (*104*) with XbaI and SpeI restriction sites and cloned into pEN_TTmiRc2 digested with SpeI and Mfe1 (Addgene #136521). CDKN1A and NRF2 PCR amplicons were prepared with SpeI and MfeI restriction sites and cloned into pEN_TTmiRc2 BirA* (Addgene #136521). Luciferase PCR amplicon was prepared with SpeI and EcoRI restriction sites and cloned into pEN_TTmiRc2 3xFLAG digested with SpeI and MfeI sites (Addgene #136519). p53DD (p53DN) and p53(R280K)-V5 PCR amplicon was prepared with SpeI and MfeI restriction sites and cloned into pEN_TTmiRc2 (Addgene #25752) digested with SpeI and MfeI (Addgene #136520, #136525).

pEN_TT donor vectors were recombined into pSLIK neo (Addgene #25735), pSLIK zeo (Addgene #25736), or pSLIK hygro (Addgene #25737) by LR recombination to obtain pSLIK 3xFLAG-Luciferase zeo (Addgene #136533), pSLIK p53DD zeo (Addgene #136534), pSLIK 3xFLAG-Luciferase hygro (Addgene #136528), pSLIK 3xFLAG-NRF2(RR) hygro (Addgene #136535), pSLIK BirA* hygro (Addgene #136537), pSLIK BirA*-CDKN1A hygro (Addgene #136538), pSLIK BirA*-NRF2 hygro (Addgene #136539), pSLIK p53(R280K)-V5 hygro (Addgene #136540) and pSLIK 3xFLAG-CHEK2 neo (Addgene #136536).

pLXSN HPV16E7 (*105*) and the ΔDLYC mutant (Addgene #136588) were provided by Scott Vande Pol. pCDH-Hyper2-puro (*64*) was provided by Joan Brugge.

### Cell lines

The MCF10A-5E clone was previously reported and cultured as described for MCF-10A cells (*13, 20*). MCF10DCIS.com cells were obtained from Wayne State University and cultured in DMEM/F-12 medium (Gibco) plus 5% horse serum (Gibco). SUM102PT cells were obtained from Asterand Biosciences and cultured in Ham’s F-12 (Gibco) plus 10 mM HEPES (Gibco), 10 ng/ml epidermal growth factor (Peprotech), 5 mM ethanolamine (Sigma), 50 nM sodium selenite (Sigma), 5 µg/ml apo-Transferrin (Sigma), 10 nM Triiodo-L-Thyronine (VWR), 5 µg/ml insulin (Sigma), 1 µg/ml hydrocortisone (Sigma), and 5% fatty acid free bovine serum albumin (VWR). SUM159PT cells were obtained from Asterand Biosciences and cultured in Ham’s F-12 (Gibco) plus 10 mM HEPES (Gibco), 5 µg/ml insulin (Sigma), 1 µg/ml hydrocortisone (Sigma), and 5% fetal bovine serum (Hyclone). All other cell lines were obtained directly from ATCC. MDA-MB-231 and MDA-MB-468 cells were cultured in L-15 medium plus 10% fetal bovine serum without supplemental CO_2_. HCC1806 and HCC1937 cells were cultured in RPMI 1640 medium plus 10% fetal bovine serum. All base media were further supplemented with 1× penicillin and streptomycin (Gibco). All cell lines are female, were grown at 37°C, authenticated by short tandem repeat profiling by ATCC, and confirmed negative for mycoplasma contamination.

### Viral transduction and selection

Lentiviruses were prepared in human embryonic kidney 293T cells (ATCC) by triple transfection of the viral vector with psPAX2 + pMD.2G (Addgene) and transduced into MCF10A-5E, MCF10DCIS.com, HCC1937, SUM159PT, MDA-MB-231, HCC1806, and MDA-MB-468 as previously described (*25*). Retroviruses were prepared similarly by double transfection of the viral vector with pCL ampho (Addgene) and transduced into MCF10A-5E cells as previously described (*22*). Transduced cells were selected in growth medium containing 2 µg/ml puromycin, 300 µg/ml G418, 100 µg/ml hygromycin, or 25 µg/ml zeocin until control plates had cleared. For RNAi-resistant addback, viral titers were adjusted to match the endogenous protein abundance as closely as possible. For mRFP1-NRF2 fluorescent reporter, we used the minimum viral titer that gave sufficient signal in sulforaphane-treated cells compared to DMSO-treated cells.

### 3D culture

3D overlay cultures were performed on top of Matrigel (BD Biosciences) as described previously for MCF-10A cells (*106*) with culture media previously optimized for each cell line (*25*). In addition, HCC1806 cells were cultured in MCF10A assay media (*106*), and SUM159PT cells were cultured in SUM159PT growth media (described above) plus 2% fetal bovine serum. For each culture, 45 µl of Matrigel was spread with a pipette tip on the bottom of an 8-well chamber slide. A suspension of 5000 single cells per well was laid on top of the Matrigel in culture media supplemented with 2% Matrigel. 3D culture medium was replaced every four days as originally described (*106*). For antioxidant supplementation, cells were treated with 50 µM Trolox (Calbiochem) for two days before 3D culture, and Trolox was included in media refeeds and supplemented every two days between refeeds. For long-term knockdown experiments, cells were treated with 1 µg/ml doxycycline (Sigma) for three days before 3D culture, and doxycycline was maintained in the 3D culture medium throughout the experiment. For experiments with long-term knockdown and inducible overexpression, cells were treated with 1 µg/ml doxycycline for two days before 3D culture, and doxycycline was maintained in the 3D culture medium throughout the experiment.

### RNA purification

RNA from cultured cells was isolated with the RNeasy Plus Mini Kit (Qiagen) according to the manufacturer’s protocol. RNA from 3D cultures at day 10 was extracted by lysing individual wells in 500 µl RNA STAT-60 (Tel-Test) and purified as described previously (*25*).

### RNA sequencing and analysis

Total RNA was diluted to 50 ng/µl and prepared using the TruSeq Stranded mRNA Library Preparation Kit (Illumina). Samples were sequenced on a NextSeq 500 instrument with NextSeq 500/550 High Output v2.5 kits (Illumina) to obtain 75-bp paired-end reads at an average depth of 15 million reads per sample. Adapters were trimmed using fastq-mcf in the EAutils package (version ea-utils.1.1.2-537) with the following options: -q 10 -t 0.01 -k 0 (quality threshold 10, 0.01% occurrence frequency, no nucleotide skew causing cycle removal). Quality checks were performed with FastQC (version 0.11.7) and multiqc (version 1.5). Datasets were aligned to the human (GRCh38.86) genome using HISAT2 with the option: --rna-strandness RF (for paired end reads generated by the TruSeq strand-specific library). Alignments were assembled into transcripts using Stringtie (version 1.3.4) with the reference guided option. Transcripts that were expressed at greater than five transcripts per million across all samples were retained for downstream analysis. Differential gene expression analysis was carried out using edgeR (version 3.8) (*107*) on raw read counts corresponding to transcripts that passed the abundance-filtering step. Trimmed Mean of M-values normalization (TMM) normalization using the calcNormFactors function was done before differential expression analysis using exactTest in edgeR. The 1,132 transcripts that were commonly differentially expressed (5% FDR) between MCF10A-5E shControl and shNRF2 #1, shControl and shNRF2 #2, and MCF10DCIS.com shControl and shNRF2 #1 are shown in Fig. 3B. Gene Set Enrichment Analysis was done on transcripts that were differentially upregulated or downregulated in shNRF2 compared to shControl using the Molecular Signatures database collections C1-C4, C6, C7 (*108, 109*). The full list of enrichments (5% FDR) is provided in file S2.

### Quantitative PCR

cDNA synthesis and quantitative PCR were performed as previously described (*25, 110*) with the primers listed in table S1. Human samples were normalized to the geometric mean of *ACTB*, *HINT1*, *PP1A*, and *TBP* (Fig. 2D and fig. S1C); *B2M*, *GAPDH*, *GUSB*, *HINT1*, *PRDX6* (fig. S1A); or *ACTB*, *B2M*, *GUSB*, *PPIA*, *PRDX6* (fig. S1B).

### Brightfield imaging and quantification of spheroid phenotypes

Brightfield 3D images were acquired on an Olympus CKX41 inverted microscope with a 4× Plan objective (four fields per chamber) and a qColor3 camera (Qimaging). Images were segmented using OrganoSeg (*111*) to produce morphometric measures for each segmented spheroid. ‘Rounded’ spheres were classified as having circularity greater than 0.9 (Fig. 3, C and E, and fig. S5, B and C). ‘Hyper-enlarged’ spheres were classified as having an area greater than e^8.5^ ~ 5000 µm^2^ (Fig. 3, D and E). ‘Proliferation suppressed’ spheres were classified as having an area less than 1600 µm^2^ for MCF10A-5E cells after 10 days of 3D culture (Fig. 2E and 4E).

### Clinical samples

Cases were identified from the pathology archives at the University of Virginia and build upon a cohort of samples previously described (*24, 25*). Hormone-negative DCIS lesions were deemed negative (less than 10% expression frequency) for estrogen receptor and progesterone receptor by clinical immunohistochemistry, and TNBC cases were additionally scored negative for HER2 amplification by clinical DNA chromogenic in situ hybridization. All clinical work was done according to IRB-HSR approval #14176 and PRC approval #1363 (502-09).

### Immunofluorescence

MCF10A-5E and MCF10DCIS.com 3D cultures were embedded at day 10 of morphogenesis, and 5-µm sections were cut and mounted on Superfrost Plus slides (Fisher). For clinical samples, paraffin tissue sections were dewaxed and antigens retrieved on a PT Link (Dako) with low-pH EnVision FLEX Target Retrieval Solution (Dako) for 20 min at 97°C. Immunofluorescence on cryosections and antigen-retrieved slides was performed as previously described (*20*) with the following primary antibodies: NRF2 (Santa Cruz #sc-13032, 1:100), phospho-Rb (Cell Signaling #8516, 1:1600), p53 (Santa Cruz #sc-126, 1:200). Slides were incubated the next day for 1 hr in the following secondary antibodies: Alexa Fluor 555-conjugated goat anti-rabbit (1:200; Invitrogen), Alexa Fluor 647-conjugated goat anti-mouse (1:200; Invitrogen).

### Image acquisition–analysis and mutual information calculation

Fluorescent images were collected on an Olympus BX51 fluorescence microscope with a 40× 1.3 numerical aperture (NA) UPlanFL oil-immersion objective and an Orca R2 CCD camera (Hamamatsu) with no binning. Images were segmented in CellProfiler (*112*) using DAPI to identify nuclei. Nuclear objects were dilated to a median diameter of 15 µm to capture approximately one whole cell. NRF2 staining was quantified in the nucleus, the whole cell, and the cytoplasm (whole cell area – nuclear area). p53 and pRB staining was quantified in the whole cell. Immunoreactivity was quantified as the median fluorescence intensity of the whole cell unless otherwise noted.

For pRB and NRF2 immunofluorescence (Fig. 1, D and E), log-transformed distributions were analyzed with the MClust function in R using the unequal variance model with either one or two mixture components specified. Model fit was evaluated by *F* test.

MCF10A-5E cells stably expressing pCDH-HyPer2-puro were imaged at 37°C in Hank’s Balanced Salt Solution (Gibco) with a 40× 1.3 NA EC Plan Neofluar oil-immersion objective on a Zeiss LSM 700 laser scanning confocal microscope. 405 nm and 488 nm lasers were used to sequentially excite two excitation peaks of HyPer-2 and collect fluorescence emission from 500– 550 nm. To calculate HyPer-2 ratios on a pixel-by-pixel basis, 488-nm images were divided by 405-nm images and thresholded in ImageJ to remove background pixel values (~10%). For quantification of cells cultured in 2D (fig. S7, B to D), the mean HyPer-2 ratio per image was used for analysis. For quantification of cells cultured as spheroids (Fig. 4C), cells were manually segmented to calculate the median HyPer-2 ratio per cell.

Clinical samples were imaged on an Olympus BX51 fluorescence microscope with a 40× 1.3 NA UPlanFL oil-immersion objective and an Orca R2 CCD camera (Hamamatsu) with 2×2 binning and fixed exposure times for NRF2 (150 msec) and p53 (50 msec). Images were autoexposed in the DAPI channel for nuclear segmentation and in the unlabeled FITC channel for autofluorescence estimation. Image fields were classified as follows: Normal—bilayered epithelium, intact basement membrane (visualized by FITC autofluorescence), and normal cytoarchitecture; DCIS—multilayered and disorganized epithelium (with partial or complete luminal filling), intact basement membrane, cytologic atypia; TNBC—invasive carcinoma cells with cytologic atypia and no discernable basement membrane. All images were segmented in CellProfiler as described above. After nuclear identification, nuclei outside of the ductal epithelium (fibroblasts, endothelial cells, and immune cells) were manually removed using the IdentifyObjectsManually module. Because paraffin fixation of tissue increases autofluorescence (*113*), the analysis excluded images that were dominated by autofluorescent bleedthrough into the Alexa 555 channel localizing NRF2. Spearman correlation was calculated between cellular FITC–555 channels and FITC–DAPI channels on a pixel-by-pixel basis for each image. Images with a FITC–555 correlation coefficient above the 95th percentile for FITC–DAPI correlation (in which autofluorescent artifacts were negligible due to the low exposure time) were excluded from further analysis.

For NRF2 quantification in neighboring cells (fig. S6), spheroid and mouse mammary gland images were loaded into CellProfiler and the IdentifyObjectManually module was used to manually identify regions of ductal epithelium. The images were cropped manually, and cell nuclei within the cropped area were identified by DAPI staining. Nuclear area was dilated to a median diameter of ~15 µm to define a cell. Position, area, and median NRF2 staining intensity were measured for each cell. Measurements were loaded into MATLAB, and single-cell NRF2 intensities were normalized to the median intensity of all exposure-matched cells. Neighboring cells were defined as cells located within a radius of 1.5 times the median cell diameter. For more distant neighbors, annular areas of 3–5 and 5–10 times the median cell diameter were used. Cells that fell within the applied search area were used to calculate the median neighbor NRF2 intensity. The original cell at the center of the search area was not included in the intensity calculations, and cells with NRF2 intensity values equal to 0 or lacking neighboring cells within the defined search area were excluded from calculations.

To quantify the association between fluorescence channels, we used mutual information in lieu of standard correlation measures (Pearson, Spearman). After appropriate transformation and binning into discrete high-low states, mutual information provides greater flexibility to capture nonlinear relationships (*114*) and more stringency to detect compressions in dynamic range (*115*). Median fluorescence intensity distributions were transformed by their respective cumulative distribution functions (probability integral transform) to produce uniformly distributed random variables (*116*). The uniform distributions were split into low and high states at the 67th percentile, and the joint–marginal state probabilities estimated for the two fluorescence channels (R, G) were used to calculate the mutual information (MI) as follows:

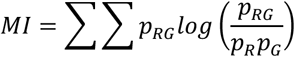

MI confidence intervals were estimated by bootstrapping the segmented cell population 1000 times. To create a randomized (null) dataset, the values of one fluorescence channel were randomly shuffled before analysis.

Clinical samples often had fewer areas of classified cells for imaging, which require an added analysis step in the mutual information calculation. For a classification (normal, DCIS, TNBC) comprised of two images from one case, we evaluated batch effects by hypergeometric test to determine if the two images separated by high-vs.-low staining intensity. If so, the case for that classification was excluded.

### Quantitative immunoblotting

Quantitative immunoblotting was performed as previously described (*25*). Primary antibodies recognizing the following proteins or epitopes were used: NRF2 (Santa Cruz Biotechnologies, #sc-13032, 1:1000), p53 (Santa Cruz Biotechnology #sc-126, 1:1000), p21 (Proteintech #10355-1-AP, 1:1000), total Chk2 (Cell Signaling #2662, 1:1000), phospho-Chk2 (Thr68, Cell Signaling #2197, 1:1000), phospho-ATM (Ser1981, Abcam #ab81292, 1:1000), KEAP1 (Santa Cruz Biotechnology #sc-15246, 1:1000), CDK4 (Cell Signaling #12790, 1:1000), CDK2 (Santa Cruz Biotechnology #sc-6248, 1:200), vinculin (Millipore #05-386, 1:10,000), GAPDH (Ambion #AM4300, 1:20,000), tubulin (Abcam #ab89984, 1:20,000), p38 (Santa Cruz Biotechnology #sc-535, 1:5000), Hsp90 (Santa Cruz Biotechnology #sc-7947, 1:5000).

### Proximity ligation using BirA*-fusions of p21 and NRF2

MCF10A-5E cells inducibly expressing the promiscuous biotin ligase BirA* (*117*), BirA*-NRF2, or BirA*-CDKN1A were plated on 10-cm plates and induced with 1 µg/ml doxycycline at 50% confluency. After 24 hours, media was refed with 1 µg/ml doxycycline, 10 µM sulforaphane (Sigma), 10 µM Nutlin-3 (Calbiochem) and 1 mM biotin (Sigma). After 24 hours, cells were lysed in 200 µl RIPA buffer (50 mM Tris (pH 8.0), 150 mM NaCl, 5 mM EDTA, 1% Triton X-100, 0.1% SDS, 0.5% sodium deoxycholate). Anti-biotin antibody enrichment of biotinylated peptides was performed as previously described (*118*). Briefly, biotin antibody bound agarose beads (ImmuneChem Pharmaceuticals Inc., #ICP0615) were washed three times in IAP buffer (50 mM MOPS (pH 7.2), 10 mM sodium phosphate and 50 mM NaCl). 500 µg (50 µl) of antibody was added to each RIPA lysate on ice. Ice-cold IAP buffer was added up to 1 ml and samples were incubated on a nutator overnight at 4°C. The next day, beads were washed four times with ice-cold IAP buffer, boiled in dithiothreitol-containing 2× Laemmli sample buffer, and used for immunoblotting against the indicated targets.

### Promoter bioinformatics

The 36 transcripts of the Fig. 1A gene cluster (*20, 24*) were assessed with four promoter analysis algorithms to identify recurrent transcription factor (TF) candidates (*119*). First, distant regulatory elements (DiRE) analysis was conducted using the DiRE website (https://dire.dcode.org) (*40*) searching evolutionary conserved 5’ untranslated regions (5’ UTR ECRs) and promoter regions (promoter ECRs) for genes on the human genome (hg18). A random set of 7500 genes was selected as background control genes. Second, Expression2Kinases (X2K) software was used to identify upstream TFs for the Fig. 1A gene cluster (*41*). The potential TFs were selected from ChIP-X Enrichment Analysis (ChEA) database using “mouse + human” as the background organisms (*120*). The *p* value from the Fisher Test and Z-score were used for sorting and ranking. Third, from the National Center for Biotechnology Information (NCBI), we collected the proximal promoter of each transcript— defined as 1416 base pairs (bp) upstream and 250 bp downstream of the transcription start site to remain within the 60 kb sequence limit—for use as an input set for MEME (*121, 122*). Using MEME-defined motifs from classic discovery mode, the top three enriched motifs were searched against the JASPAR CORE (2018) database (containing 1404 defined TF binding sites for eukaryotes) (*123*) or HOCOMOCO Human (v11) database (containing 769 TF binding motifs) (*124*) using TOMTOM (*125*) to identify transcription factor recognition sequences. Last, oPOSSUM (*43*) was used to identify potential TFs targeting transcripts in the cluster. We selected Single Site Analysis – Human mode and used all 24,752 genes in the oPOSSUM database as a background. All vertebrate profiles with a minimum specificity of 8 bits in the JASPAR CORE Profiles were selected as TF binding sites sources. oPOSSUM was run with the following parameters: conservation cutoff of 0.4, matrix score threshold of 85%, amount of upstream/downstream sequence: 2000/0, and sort results by Fisher score.

### Computational modeling

The NRF2 pathway was encoded as first- and second-order rate equations for KEAP1 oxidation and NRF2 stabilization; NRF2-mediated transcription of antioxidant enzymes was modeled as a Hill function (*66, 67*). The p53 pathway was reconstructed from a delay differential equation model of p53 signaling in response to DNA damage (*69*). Abundances in original p53 model were unitless, but abundances were cast as concentrations in the earlier NRF2 models. Consequently, the integrated model adopted unitless abundances in its initial conditions and second-order parameters (table S2). To adapt the p53 DNA-damage model to respond to oxidative stress, we changed the ‘Signal’ activation (representing activation of upstream kinases p-ATM and p-CHEK2) from a Heaviside step function to a first-order oxidation reaction of ATM/CHEK2 by intracellular H_2_O_2_ (*61*). A basal ROS generation rate was added yielding a realistic intracellular H_2_O_2_ burden at steady state (*68*). Transcription of antioxidant enzymes by p53 (*70*) was modeled using the same model parameters describing the p53-mediated induction of MDM2 (*69*). p53- and NRF2-mediated antioxidant gene transcription contribute to a shared pool of antioxidant enzymes, which catalytically reverse the oxidation states of KEAP1 and p-ATM/CHEK2. Transcription of CDKN1A by p53 (*126*) was included for model calibration (fig. S10) and for testing the relevance of p53–p21–NRF2 crosstalk (see below). The integrated base model of NRF2–p53 oxidative-stress signaling contains 42 reactions and 22 ordinary differential equations (ODEs). The model was simulated with dde23 in MATLAB to reach steady state before the addition of oxidative stress.

The integrated model was calibrated to capture the dynamics of MCF10A-5E cells stimulated with 200 µM H_2_O_2_ (fig. S10). Bolus addition of H_2_O_2_ was simulated as an impulse of intracellular H_2_O_2_. We used an H_2_O_2_ partition coefficient that gave rise to NRF2 stabilization levels comparable to immunoblot quantification (extracellular / intracellular partition = 3). We approximated p-ATM/CHEK2 in the integrated model as the maximum normalized increase of p-ATM or p-CHEK2 over baseline at each experimental time point. Robustness of the system output to initial conditions was evaluated by randomly varying the concentration of model species with a log coefficient of variation of 10% taking the base model as the geometric mean.

For simulations involving the mRFP1-NRF2 reporter (NRF2rep, Fig. 5C), NRF2rep and mature fluorescent species (Nrf2repmat) were added to the MCF10A-5E base model. Both reporter species were allowed to react with KEAP1, but neither could bind Maf proteins or antioxidant response elements in the model (fig. S7E). We used an mRFP1 maturation time of one hour (*74*) to model the conversion of NRF2rep to NRF2repmat. The modifications added 19 additional reactions and eight additional ODEs to the MCF10A-5E base model.

For simulations involving p53–p21–NRF2 crosstalk (fig. S10C), we added reactions involving p21 binding to NRF2 to the MCF10A-5E base model. p21 was assumed to interact with NRF2 like KEAP1 and compete with KEAP1 for binding NRF2 through its DLG and ETGE domains (*71*). The p21:NRF2 complex was assumed to degrade at the same reduced rate as when NRF2 is bound to oxidized KEAP1 (k_nrf2degox). These modifications added eight additional reactions and two additional ODEs to the MCF10A-5E base model.

For simulations involving MCF10DCIS.com cells (Fig. 5, E to G), RNA-sequencing data (Fig. 3B) was used to estimate proportional differences in model species abundance between MCF10DCIS.com and MCF10A-5E cells. Average gene expression in transcripts per million (TPM) from the four biological replicates of MCF10DCIS.com and MCF10A-5E control cell lines was calculated for each gene. Fold changes in model species of MCF10DCIS.com relative to MCF10A-5E were used to adjust each initial condition in the model. For the ‘MAF’ species, we used the median fold change in NRF2-binding small Mafs *MAFF*, *MAFG*, and *MAFK* (*127*). For the antioxidant species, we used the median fold-change in *TXN*, *SOD1*, *PRDX1*, and *HMOX1* to include antioxidants that react with both free radicals and oxidized proteins (*128*). Additionally, the MCF10DCIS.com model included a 1.4-fold increase in the basal ROS generation rate, informed by the increased median HyPer2-ratio in MCF10DCIS.com cells compared to MCF10A-5E cells (fig. S7D). The increased ROS generation rate was paired with an increase basal turnover of the antioxidant pool to arrive at steady-state antioxidant gene expression levels consistent with MCF10DCIS.com RNA-seq data.

For simulations involving bursts of oxidative stress, an increased ROS production rate was added for two hours to match the duration of transient stabilizations of JUND (a gene in the NRF2-associated gene cluster) in 3D (*24*). We selected the minimum increase in ROS generation that gave rise to a detectable stabilization of both the NRF2 and p53 pathways in the MCF10A-5E base model. For MCF10DCIS.com and TNBC models, the mean ROS generation rate was scaled 1.4-fold to reflect the increased basal ROS generation rate described above. NRF2 knockdown was encoded by decreasing the net synthesis rate of NRF2 fivefold to mimic the fivefold decrease in NRF2 protein resulting from short-hairpin knockdown (Fig. 2C). To account for secondary transcriptional adaptations (Fig. 5F), initial conditions were also adjusted by RNA-seq-based fold changes in model species for shNRF2 cells relative to negative-control cells (Fig. 3B). Dominant negative p53 was encoded by removing all reactions downstream of p53 (transcriptional activation of *MDM2*, *PPM1D*, *CDKN1A*, and the p53 share of the antioxidant enzyme pool).

For the control case and all genetic perturbations (shNRF2, DNp53, and shNRF2+DNp53), 100 simulations were run with random ROS generation rates varied with a log coefficient of variation of 25% to capture the variability of HyPer-2 ratios observed experimentally (Fig. 4D). Each simulation was run for two hours with increased ROS production rate and then an additional 10 hours to allow relaxation back to steady state. For assessment of species coordination (Fig. 5, C to E, and 7B), species abundances were captured at 10 random timepoints from each simulation and mutual information was calculated as it was for quantitative immunofluorescence datasets. For oxidative stress analysis (Fig. 5, F and G, and 7B), the time-integrated intracellular H_2_O_2_ concentration was used as an overall measure of oxidative stress.

For simulations involving TNBC cells (Fig. 7, A and B), RNA-seq data from the NIH LINCS consortium (*86*) (HMS dataset ID: 20348) was used to estimate proportional differences in model species abundance between 15 TNBC cell lines and MCF10A cells. Reads per kilobase per million mapped reads values were normalized as transcripts per million before fold-change calculation. MAF and antioxidant species were estimated as described above. TNBC models used the same increased basal ROS generation rate as in the MCF10DCIS.com model (*129*). To simulate p53 mutation in the 15 p53-mutant TNBC cell lines, all reactions downstream of p53 were removed.

### Statistical analysis

For analysis of the 10cRNA-seq dataset (Fig. 2A), Spearman correlation between transcripts and the median expression of the NRF2-associated gene cluster was calculated at a false-discovery rate of 10%. Transcripts with a Spearman correlation coefficient above 0.5 were examined by Gene Ontology analysis. Statistical enrichment of Gene Ontology terms was assessed by Fisher exact test with FDR-corrected *p* values. For quantitative PCR data, differences in geometric means were assessed by Welch’s *t* test after log transformation (fig. S1, A and B). Statistical interaction between shNRF2 and DNp53 and differences between immunoblotting time courses were assessed by two-way ANOVA (Fig. 2, D and E, 3E, 4, A and B, 5, F and G, and fig. S1C). Statistical interaction between shNRF2, DNp53, and Trolox was assessed by three-way ANOVA (Fig. 4E). For fig. S11B, two-way ANOVA without replication was used. For unpaired clinical data, multi-group comparison was made by Kruskal-Wallis rank-sum test (Fig. 6, D to F). For 3D spheroid growth, mean differences in area were assessed by Kruskal-Wallis test with Dunn’s post-hoc test (Fig. 7, C to G). Distributions were compared by Kolmogorov-Smirnov test (fig. S1D, S7, and S9). All other two-sample comparisons were performed by Student’s *t* test.

## Acknowledgments

We thank Page Murray for assisting with quantitative PCR, Pat Pramoonjago for help with processing the clinical samples, Emily Farber at the Center for Public Health Genomics for performing the RNA sequencing, Scott Vande Pol and Joan Brugge for plasmid reagents, and Amy Bouton and Janet Cross for critically reading this manuscript.

## Author contributions

Unless otherwise noted, E.J.P. performed the experiments, modeling, and bioinformatic analyses in the manuscript and drafted the manuscript. J.S.B. performed quantitative PCR, led the image analysis of local NRF2 stabilization, and contributed experiments related to the NRF2–p53 computational model. C.Y.L. built the initial draft of the NRF2–p53 computational model and contributed experiments related to model construction. T.M. assisted with immunocytochemistry and imaging. D.C. and L.W. provided cloning assistance. K.A.A. identified the clinical samples, performed the p53 immunohistochemical scoring, and reviewed the pathology images in the manuscript. C.C.W. completed the promoter bioinformatics along with the initial genetic perturbations and phenotyping related to the work. K.A.J. assisted with image acquisition for the clinical samples, edited the final manuscript, supervised all research, and secured funding for the work.

## Data and materials availability

RNA sequencing data are available through the NCBI Gene Expression Omnibus (GSE141228, https://www.ncbi.nlm.nih.gov/geo/query/acc.cgi?acc=GSE141228 Reviewer token: ozqzgqewhturpsf). Plasmids related to this work are deposited with Addgene (#136519–136528, #136533–136540, #136579–136581, #136583–136585, #136587–136588).

## Supplementary Materials

**Fig. S1.**
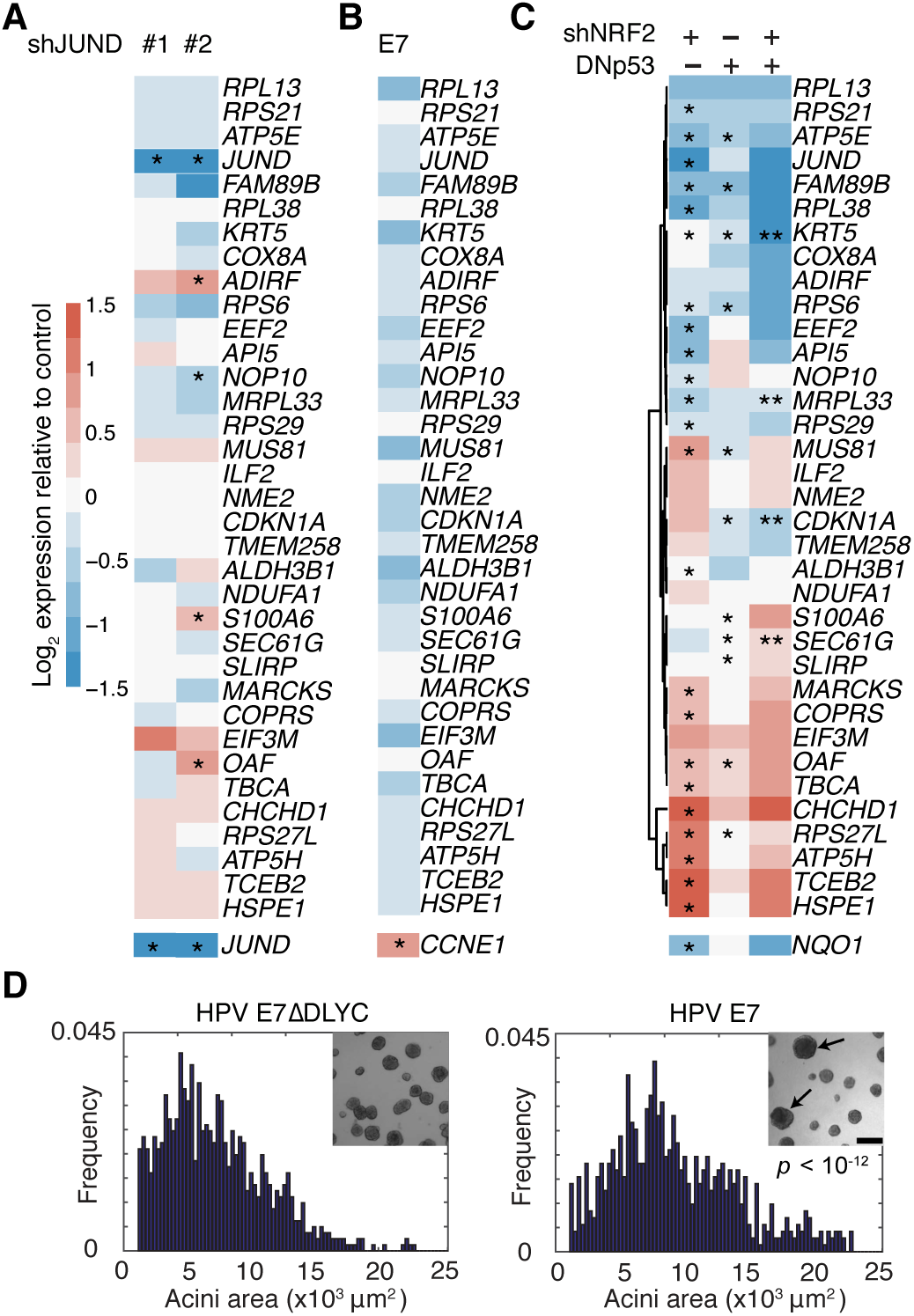
Abundance of the heterogeneously regulated gene cluster is perturbed by NRF2 knockdown or p53 disruption but not by JUND knockdown or human papillomavirus (HPV) E7-induced inhibition of RB. (**A**) Inducible JUND knockdown does not reliably affect transcripts within the gene cluster other than *JUND*. (**B**) Constitutive inhibition of RB with HPV E7 does not significantly affect transcript abundance within the gene cluster. The E2F1 target gene *CCNE1* was used as a control for efficacy of ectopic E7 expression. (**C**) Single and combined perturbations of NRF2 and p53 have complex effects on the gene cluster. The NRF2 target gene *NQO1* was used as a control for efficacy of shNRF2. (**D**) E7 expression elicits hyperproliferation (*23*) compared to a ΔDLYC control that does not bind RB (*130*). For (A) and (C), MCF10A-5E cells with or without JUND or NRF2 knockdown or DNp53 were treated with 1 µg/ml doxycycline for 48 hr, grown as 3D spheroids for 10 days, and profiled for the indicated genes by quantitative PCR. For (B) and (D), MCF10A-5E cells stably expressing E7 or E7ΔDLYC were grown as 3D spheroids for 10 days and profiled for the indicated genes by quantitative PCR or imaged by brightfield microscopy and segmented after 22 days. For (A) to (C), data are shown as the log_2_ geometric mean relative to the negative control (shGFP (A and C) or E7ΔDLYC (B) with or without FLAG-tagged LacZ (C)), with asterisks indicating significant changes or interaction effects (rightmost column of (C)) by two-way ANOVA of *n* = 8 (A and C) or 4 (B) independent 3D-cultured samples and a false-discovery rate of 5%. For (D), size histograms were compared by K-S test and scale bars are 200 µm.

**Fig. S2.**
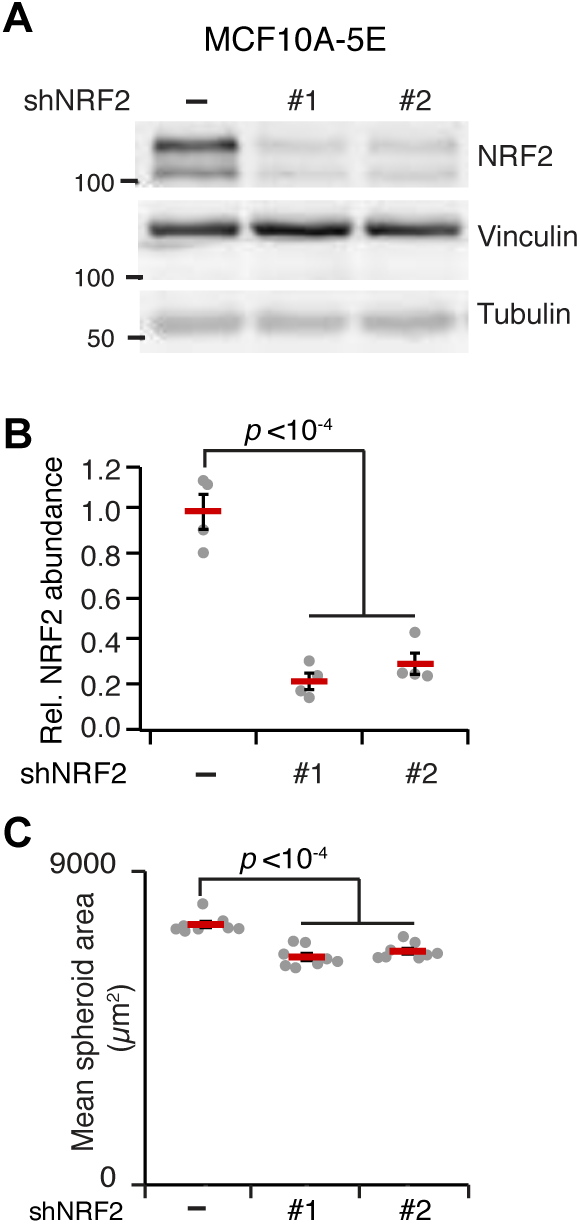
NRF2 knockdown and 3D phenotype quantification in MCF10A-5E cells. (**A**) Knockdown of endogenous NRF2 by shRNA in doxycycline-treated MCF10A-5E cells. MCF10A-5E cells were treated with 1 µg/ml doxycycline for 72 hr and immunoblotted for NRF2 with vinculin and tubulin used as loading controls. (**B**) Densitometry from replicated NRF2 knockdown. Data are shown as the mean ± s.e.m. of *n* = 4 biological replicates. (**C**) Inducible knockdown of NRF2 causes mild growth inhibition of MCF10A-5E 3D spheroids. Data are shown as the mean ± s.e.m. of *n* = 8 independent 3D-cultured samples after 10 days.

**Fig. S3.**
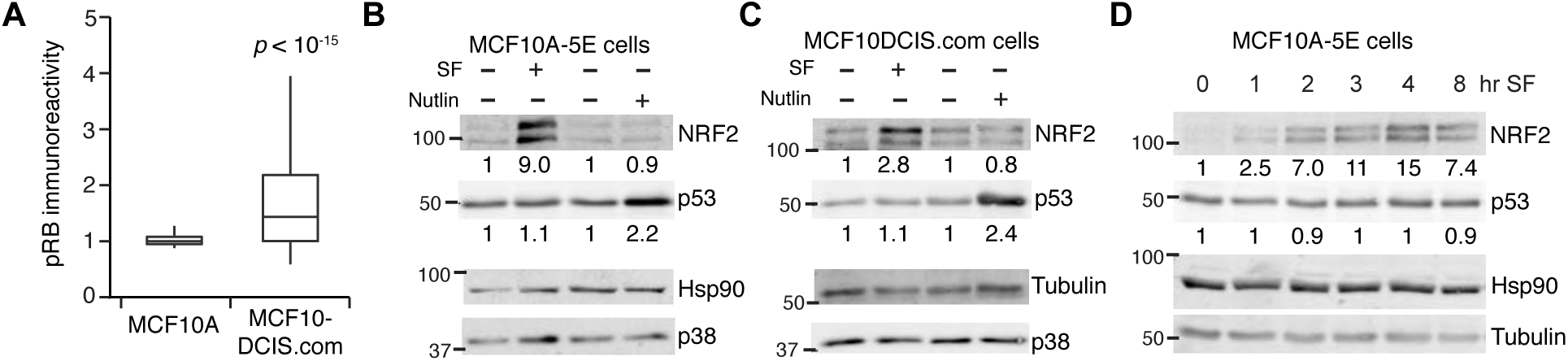
Proliferation differences and signaling similarities between MCF10A-5E and MCF10DCIS.com cells. (**A**) MCF10DCIS.com cells show sustained RB phosphorylation (pRB) in 3D cultures compared to MCF10A-5E spheroids. pRB staining was quantified after 10 days of 3D culture collected from *n* = 1350 cells from 60–70 spheroids per cell line. (**B** and **C**) Both MCF10A-5E and MCF10DCIS.com cells stabilize NRF2 in response to the electrophile sulforaphane (SF) and stabilize p53 in response to the MDM2 inhibitor Nutlin-3. **(D)** p53 is not stabilized by SF for up to eight hours in MCF10A-5E cells. Cells were treated with 10 µM SF for two hours or 10 µM Nutlin-3 for four hours (B and C) or 10 µM SF for the indicated times (C) and immunoblotted for NRF2 and p53 with Hsp90, tubulin, and p38 used as loading controls. Representative immunoblots are shown from biological duplicates.

**Fig. S4.**
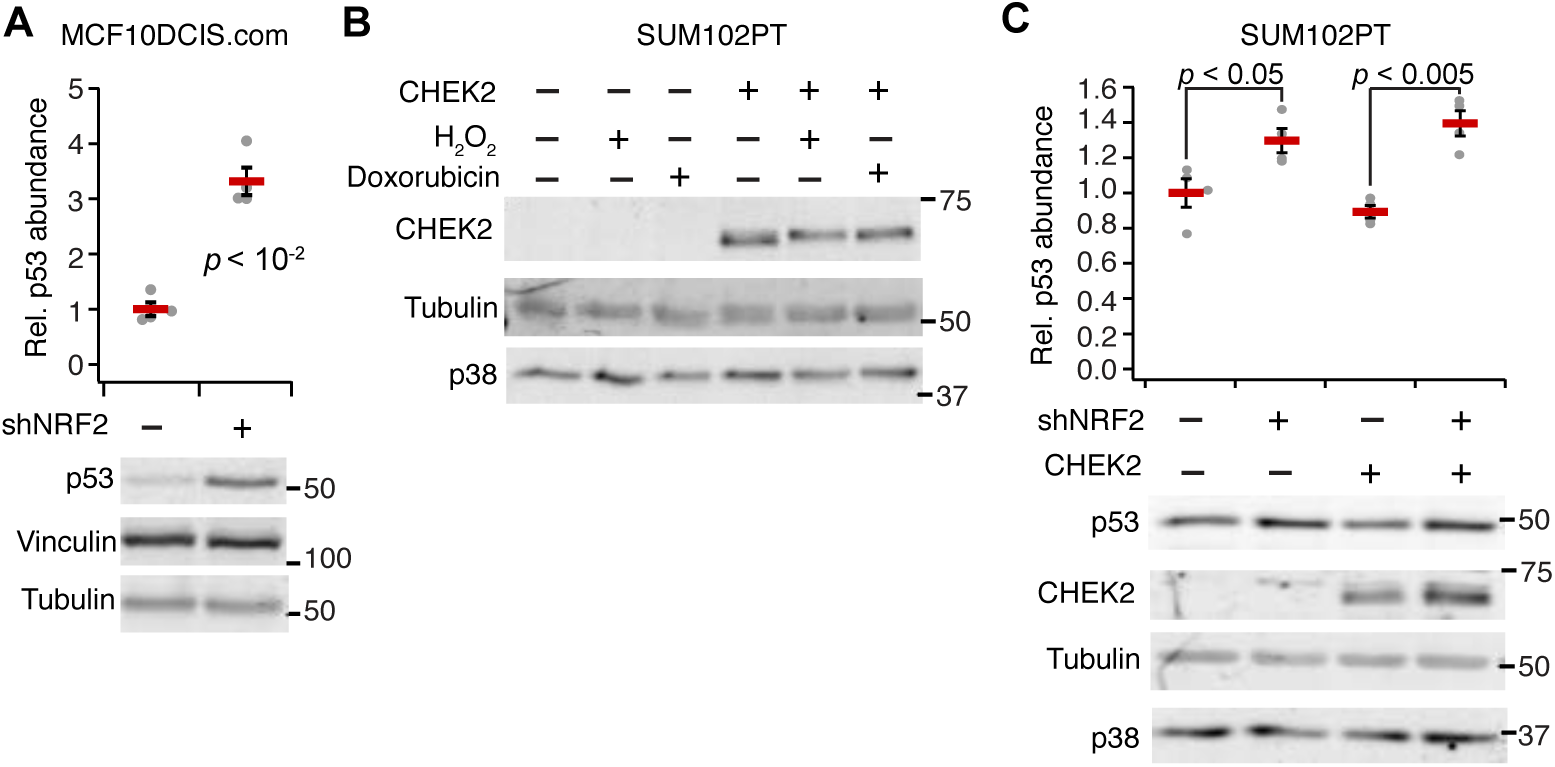
NRF2 knockdown causes p53 stabilization in premalignant breast epithelial cell lines. (**A**) Inducible knockdown of NRF2 causes p53 stabilization in MCF10DCIS.com cells. MCF10DCIS.com cells were treated with 1 µg/ml doxycycline for 72 hr and immunoblotted for p53 with vinculin and tubulin used as loading controls. Representative immunoblots are shown of *n* = 4 biological replicates. (**B**) Reconstituted CHEK2 expression in *CHEK2^1100delC^* SUM102PT cells. Note that reconstituted CHEK2 is upshifted upon H_2_O_2_-induced oxidative stress and doxorubicin-induced DNA damage, suggesting modification by upstream kinases. (**C**) Inducible knockdown of NRF2 causes p53 stabilization in SUM102PT cells and becomes more pronounced upon reconstitution of wild-type CHEK2. SUM102PT cells were treated with 1 µg/mL doxycycline for 72 hr and immunoblotted for p53 and CHEK2 with tubulin and p38 used as loading controls. Representative immunoblots are shown as the mean ± s.e.m. of *n* = 4 biological replicates.

**Fig. S5.**
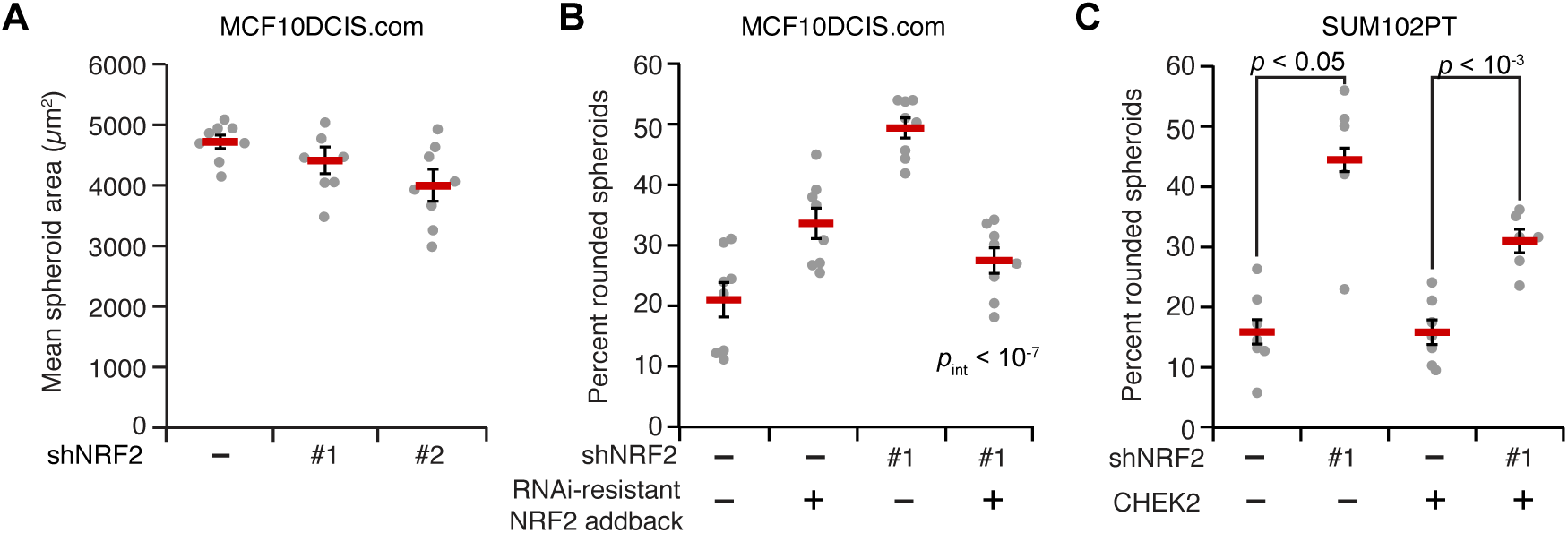
Premalignant breast epithelial cell lines have similar adaptations to NRF2 knockdown in spheroid culture. (**A**) NRF2 knockdown in MCF10DCIS.com cells does not alter spheroid growth. (**B**) NRF2 knockdown in MCF10DCIS.com cells causes an increase in rounded spheroids (Fig. 3C) that is reverted upon addback of an RNAi-resistant version of NRF2. (**C**) NRF2 knockdown in SUM102PT cells causes increased rounding with or without wild-type CHEK2 reconstitution. Data are shown as the mean ± s.e.m. of *n* = 8 independent 3D-cultured samples after 8 days (A and B) and *n* = 6 independent 3D-cultured samples after 16 days (C). For (C), cultures were treated with 1 µg/ml doxycycline at day 8 and analyzed at day 16. For (B) and (C), rounded spheroids were gated as in Fig. 3, C and E.

**Fig. S6.**
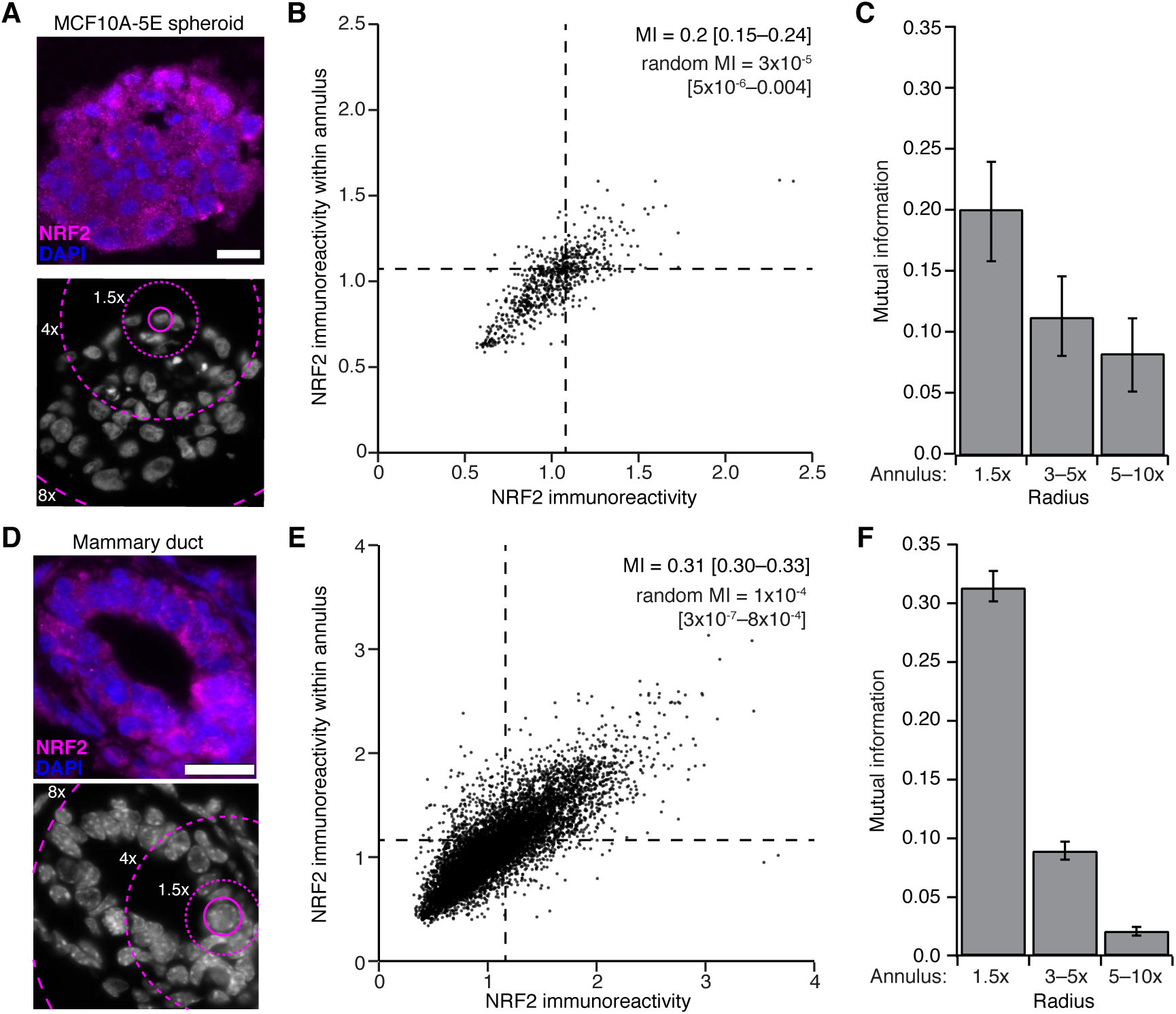
Local niches of NRF2 stabilization in MCF10A-5E 3D spheroids and pubertal murine mammary glands. (**A** to **C**) Quantification of neighboring NRF2 stabilization in MCF10A-5E 3D spheroids. (**D** to **F**) Quantification of neighboring NRF2 stabilization in murine mammary glands during puberty. In (A) and (D), representative merged images for NRF2 (magenta) with DAPI nuclear counterstain (blue) are shown (top). DAPI stain is shown below with magenta rings defining neighboring annuli used for mutual information (MI) calculations. In (B) and (E), MI is shown between NRF2 staining in single cells and surrounding cells within a radius equal to 1.5x the median cell diameter for *n* = 737 MCF10A-5E cells and 10316 mammary epithelial cells. In (C) and (E), MI is shown between NRF2 staining in single cells and surrounding cells that fall within annuli of different radius from a single cell. MI is shown with 90% CI estimated from *n* = 1000 bootstrap replicates.

**Fig. S7.**
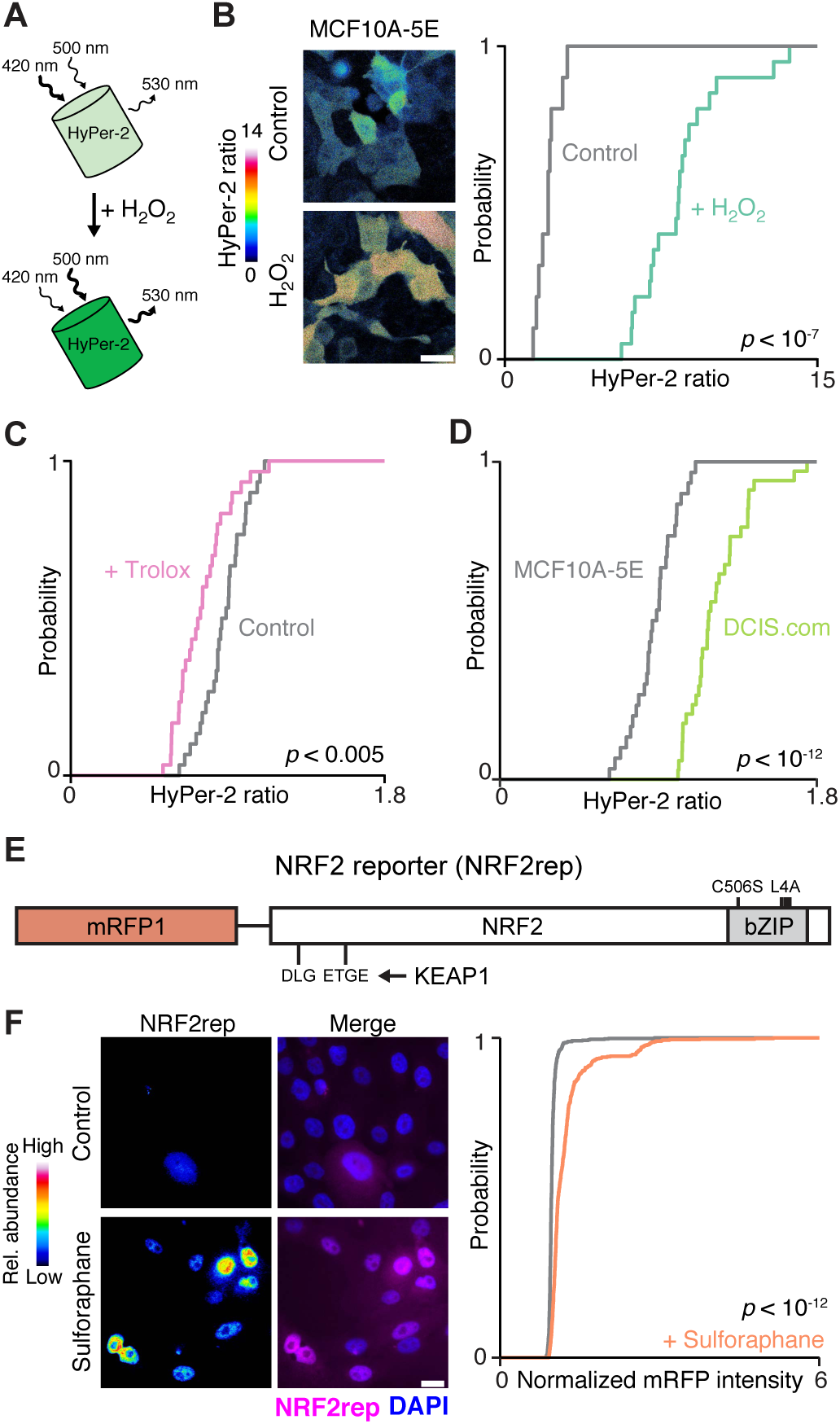
Description and validation of the HyPer-2 probe for H_2_O_2_ and the mRFP1-NRF2 reporter. (**A**) Schematic of HyPer-2. Increases in intracellular H_2_O_2_ cause an increase in 530 nm fluorescence upon excitation at 500 nm and a reduction upon excitation at 420 nm. (**B**) HyPer-2 fluorescence ratios increase in MCF10A-5E cells treated with 200 µM H_2_O_2_. (**C**) HyPer-2 fluorescence ratios are decreased in MCF10A-5E cells cultured with the antioxidant Trolox (50 µM for 48 hours). (**D**) HyPer-2 fluorescence ratios are significantly elevated in transformed MCF10DCIS.com (DCIS.com) cells compared to MCF10A-5E cells. (**E**) Schematic of the mRFP1-NRF2 reporter (NRF2rep). NRF2 is fused to mRFP1 at its N-terminus, and the DNA-binding domain is mutated (C506S) along with four leucines (L4A) in the leucine zipper region of the bZIP domain. NRF2rep remains targeted by KEAP1 through its DLG and ETGE binding motifs. (**F**) NRF2rep increases in MCF10A-5E cells treated with 10 µM sulforaphane for 2 hours. Representative pseudocolored images for NRF2rep (left) are shown merged with DAPI nuclear counterstain (right). For (B) to (D) and (F), HyPer-2 fluorescence ratios and mRFP1-NRF2 fluorescence are summarized as cumulative density plots. Scale bars are 20 µm.

**Fig. S8.**
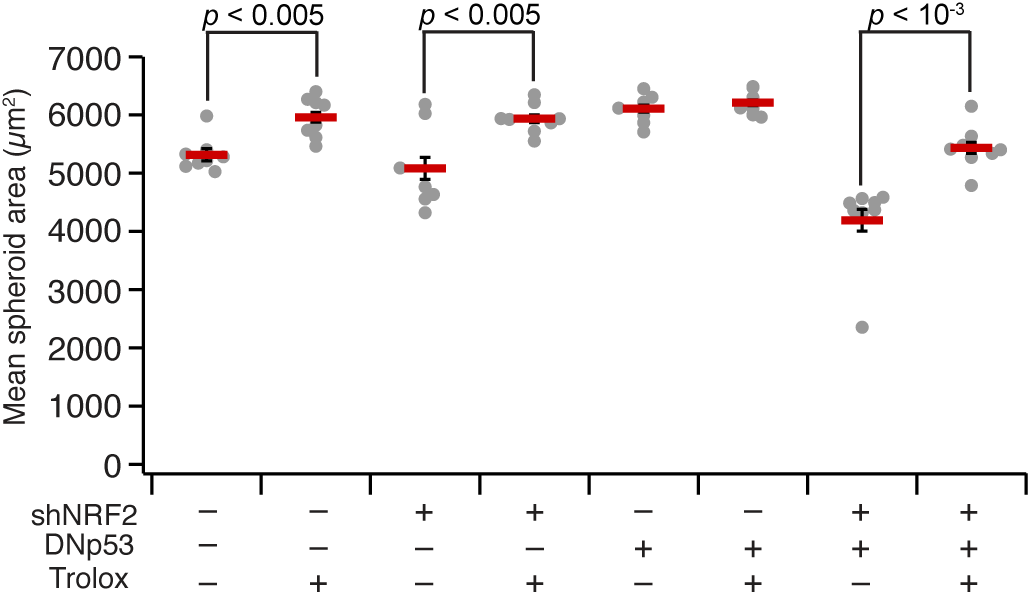
Antioxidant treatment causes an overall increase in MCF10A-5E spheroid size. MCF10A-5E cells were treated with 50 µM Trolox for two days before 3D culture, and Trolox was included in media refeeds and supplemented every two days between refeeds. Data are shown as the mean ± s.e.m. of *n* = 8 independent 3D-cultured samples after 10 days.

**Fig. S9.**
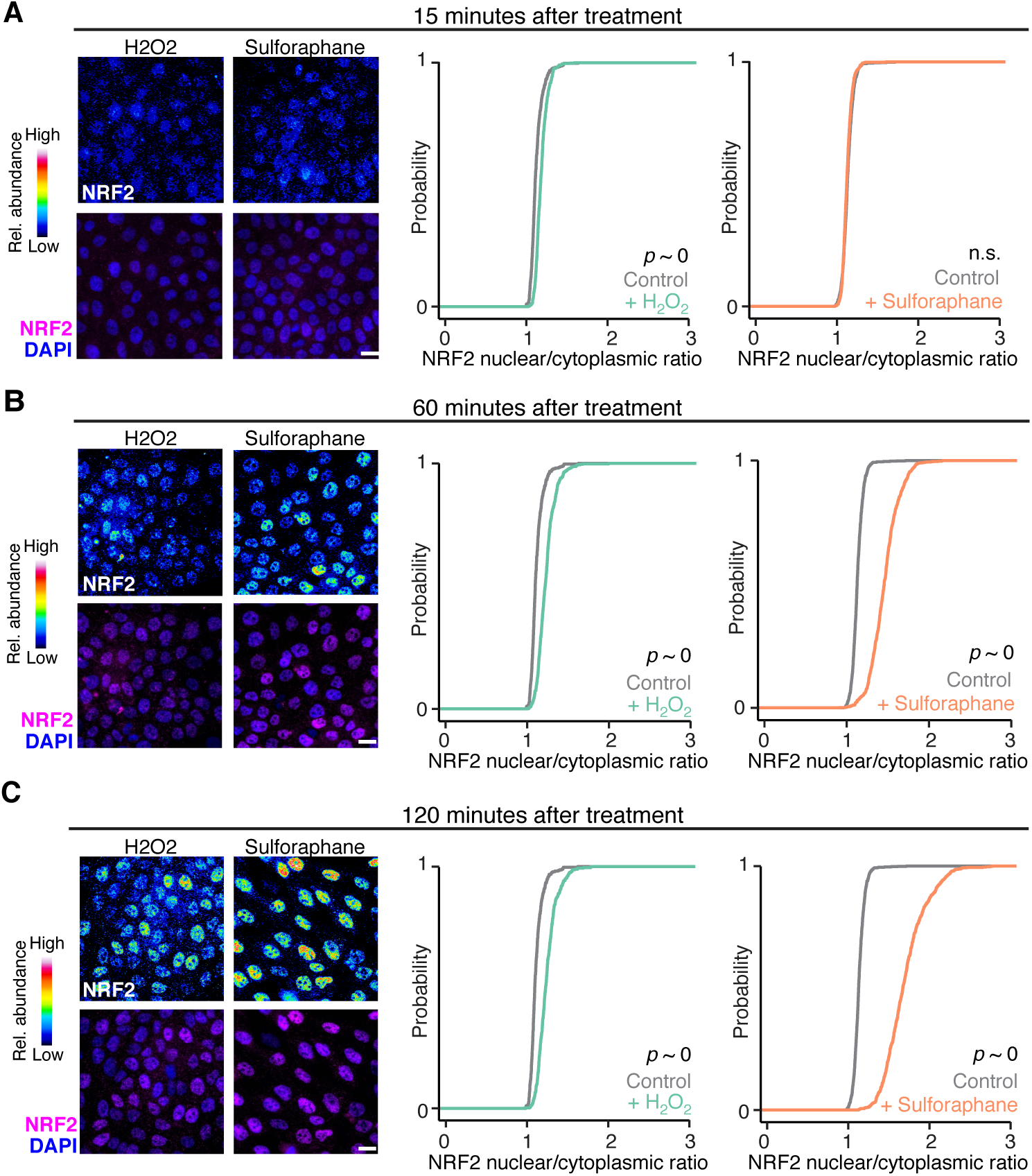
Oxidative stress stabilizes NRF2 in the cytoplasm more so than electrophilic stress. (**A** to **C**) Time course of NRF2 stabilization in MCF10A-5E cells were treated with 200 µM H_2_O_2_ or 10 µM sulforaphane for the indicated times and analyzed by immunofluorescence for NRF2 from *n* = 400–800 cells per time point. For each subpanel, representative pseudocolored images for NRF2 are shown on the top and merged with DAPI nuclear counterstain on the bottom. Scale bar is 20 µm. Cytoplasmic and nuclear NRF2 immunoreactivity is summarized as cumulative density plots.

**Fig. S10.**
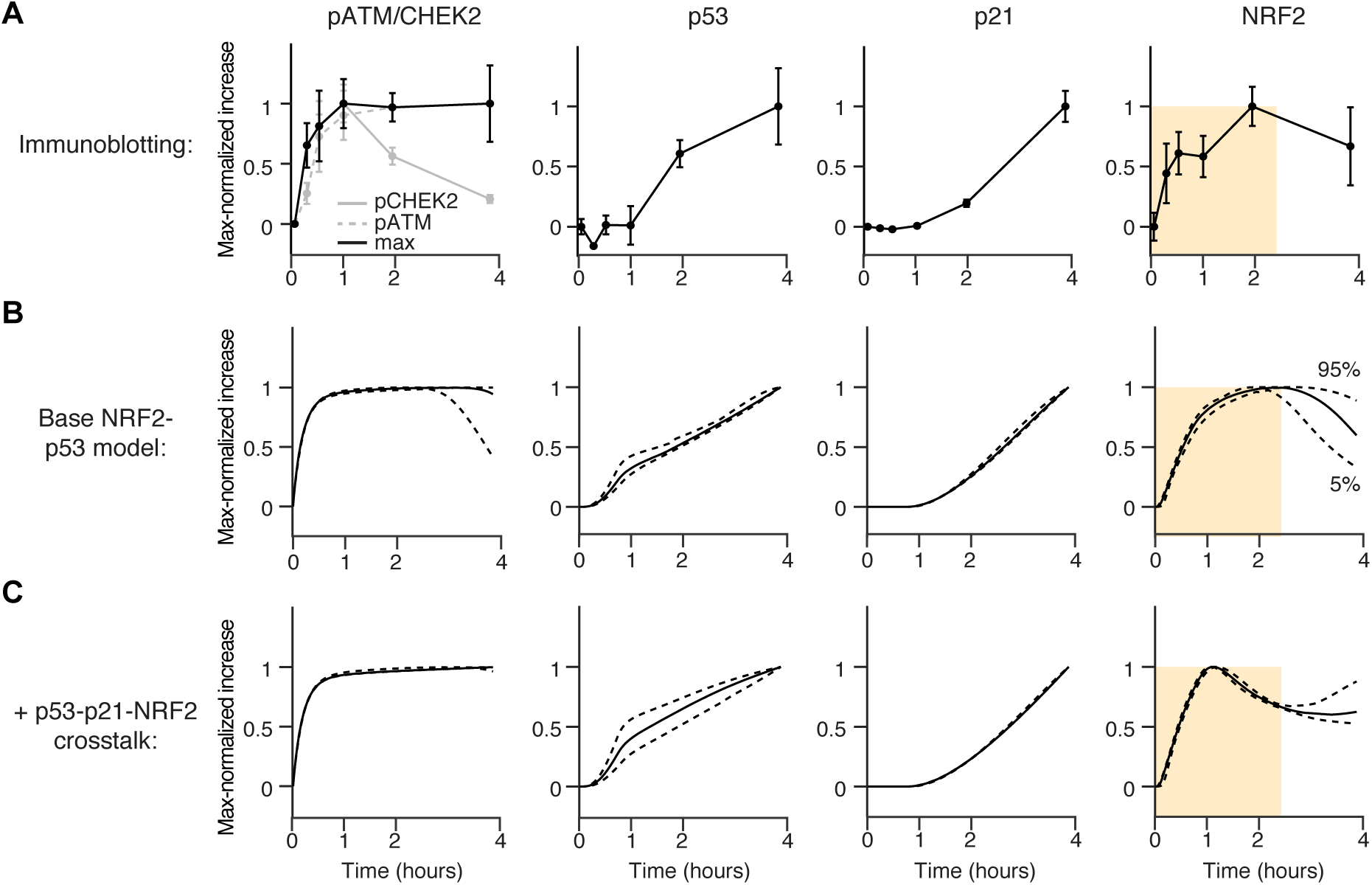
Calibration of an integrated NRF2–p53 systems model for oxidative stress. MCF10A-5E cells were treated with 200 µM H_2_O_2_ for the indicated times. (**A**) Quantitative immunoblotting for phospho-CHEK2 (pCHEK2 Thr68), phospho-ATM (pATM Ser1981), total p53, total p21, and total NRF2. Data are shown as the mean ± s.e.m. of *n* = 4 biological replicates. The maximum of pCHEK2 or pATM (max) was taken as the pATM/CHEK2 value for model calibration. (**B**) Calibration of the integrated base model to experimental data. (**C**) Addition of p53–p21–NRF2 crosstalk to the base model in (B). For (B) and (C), model results are shown as the median (solid) ± 90% confidence interval (dashed) from *n* = 50 simulations of initial conditions varied with a lognormal distribution of 10% about the geometric mean. Experiments and simulations are shown as the max-normalized increase above baseline, which was set to zero. The shaded region indicates the kinetic discrepancies between the two models relative to experiments.

**Fig. S11.**
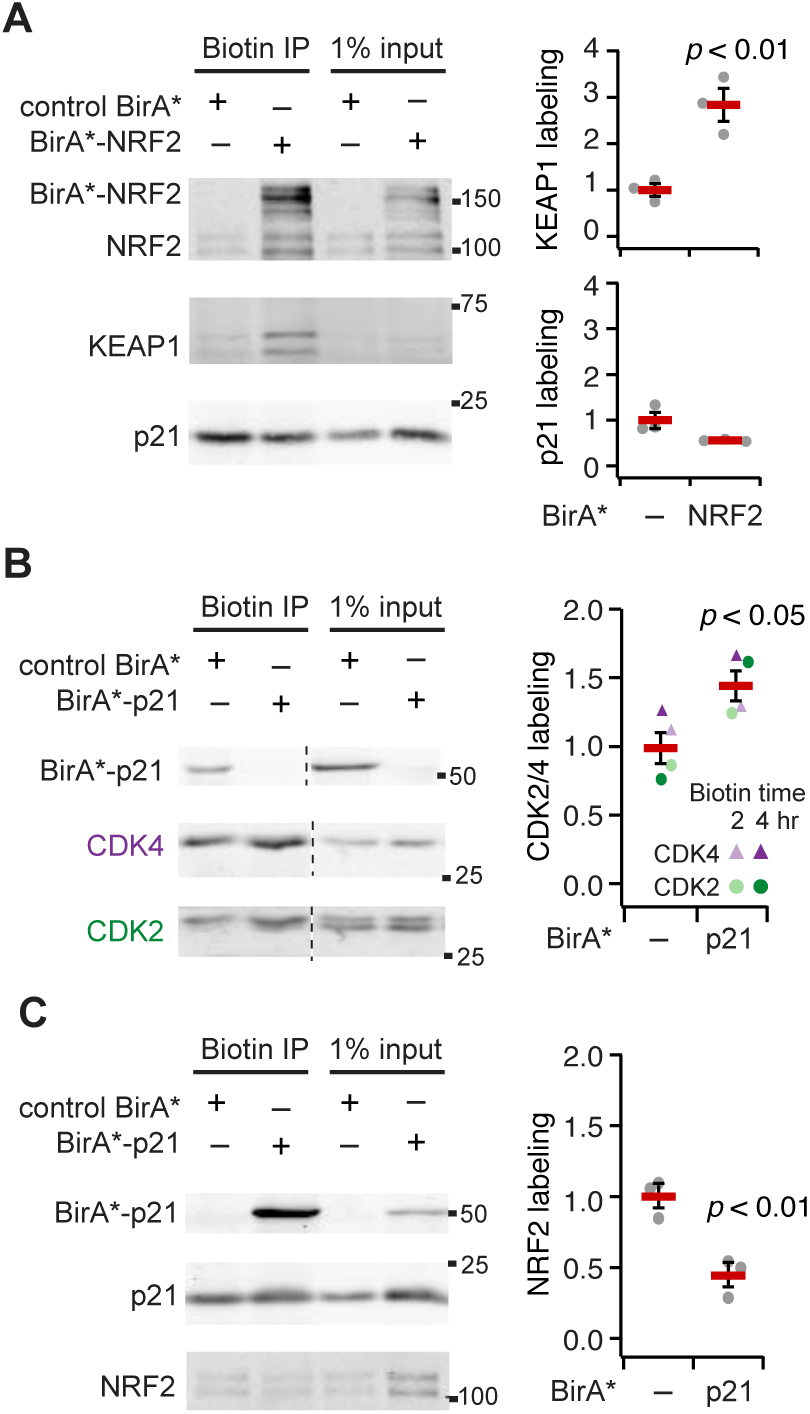
Endogenous NRF2 and p21 are not proximity labeled by BirA* fusions of each other. (**A**) BirA*-NRF2 labels endogenous KEAP1 but not p21. (**B** and **C**) BirA*-p21 labels endogenous CDK4 and CDK2 (B) but not NRF2 (C). MCF10A-5E cells were treated with 1 µg/ml doxycycline for 48 hours, 10 µM sulforaphane, 10 µM Nutlin-3 and 1 mM biotin for 24 hours (A and B) or 2 and 4 hours as indicated (C). Dashed lines indicate noncontiguous lanes on the same immunoblot. Input and biotin immunoprecipitated lysates were immunoblotted for NRF2, p21, KEAP1, CDK4, and CDK2. Representative immunoblots are shown for *n* = 3 biological replicates for (A) and (C).

**Fig. S12.**
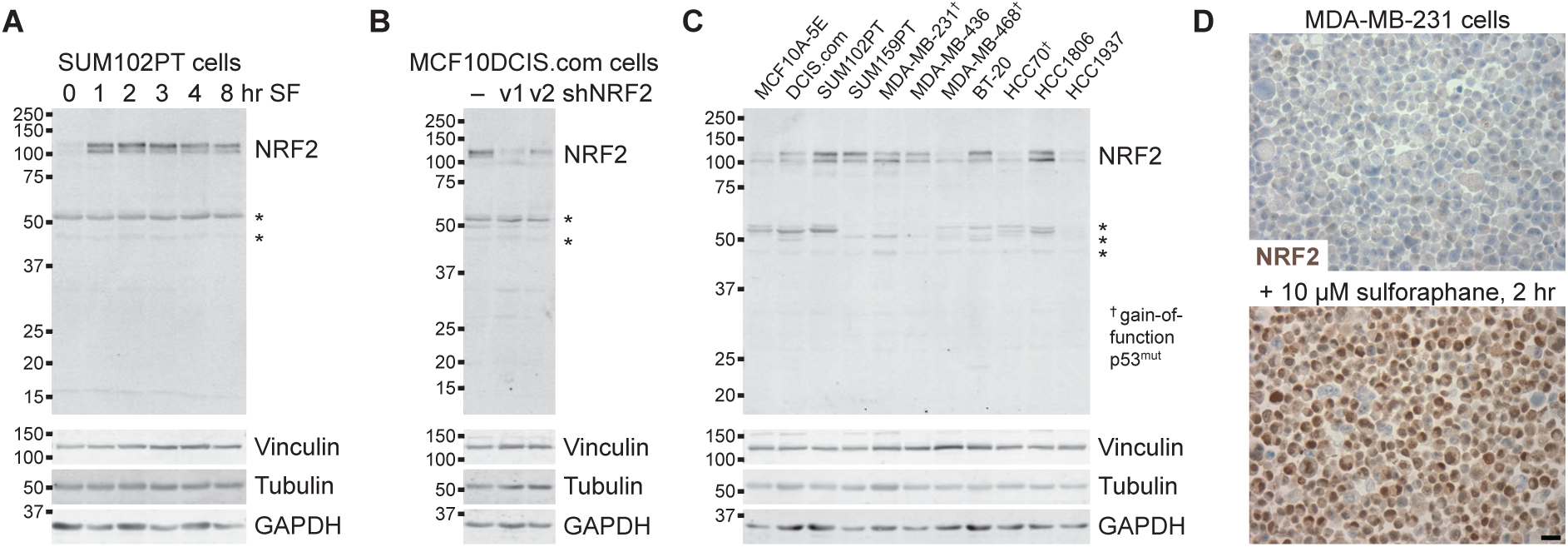
Anti-NRF2 antibody validation for immunohistochemistry. (**A**) NRF2 is the predominant band induced with the electrophile sulforaphane (SF) in SUM102PT cells. Similar results were obtained with various cell lines. (**B**) Specificity of the ~100 kDa NRF2 immunoreactive band confirmed by shRNA-mediated knockdown in MCF10DCIS.com cells. Similar results were obtained with various cell lines. (**C**) NRF2 abundance varies across triple-negative cell lines. Breast cancer lines with gain-of-function p53 are indicated (†). (**D**) Increased immunohistochemical staining in MDA-MB-231 cell pellets treated with sulforaphane as indicated. Scale bar is 20 µm. For (A) to (C), asterisks indicate minor nonspecific bands that are likely denaturation-induced epitopes based on the staining of control cell pellets in (D).

**Fig. S13.**
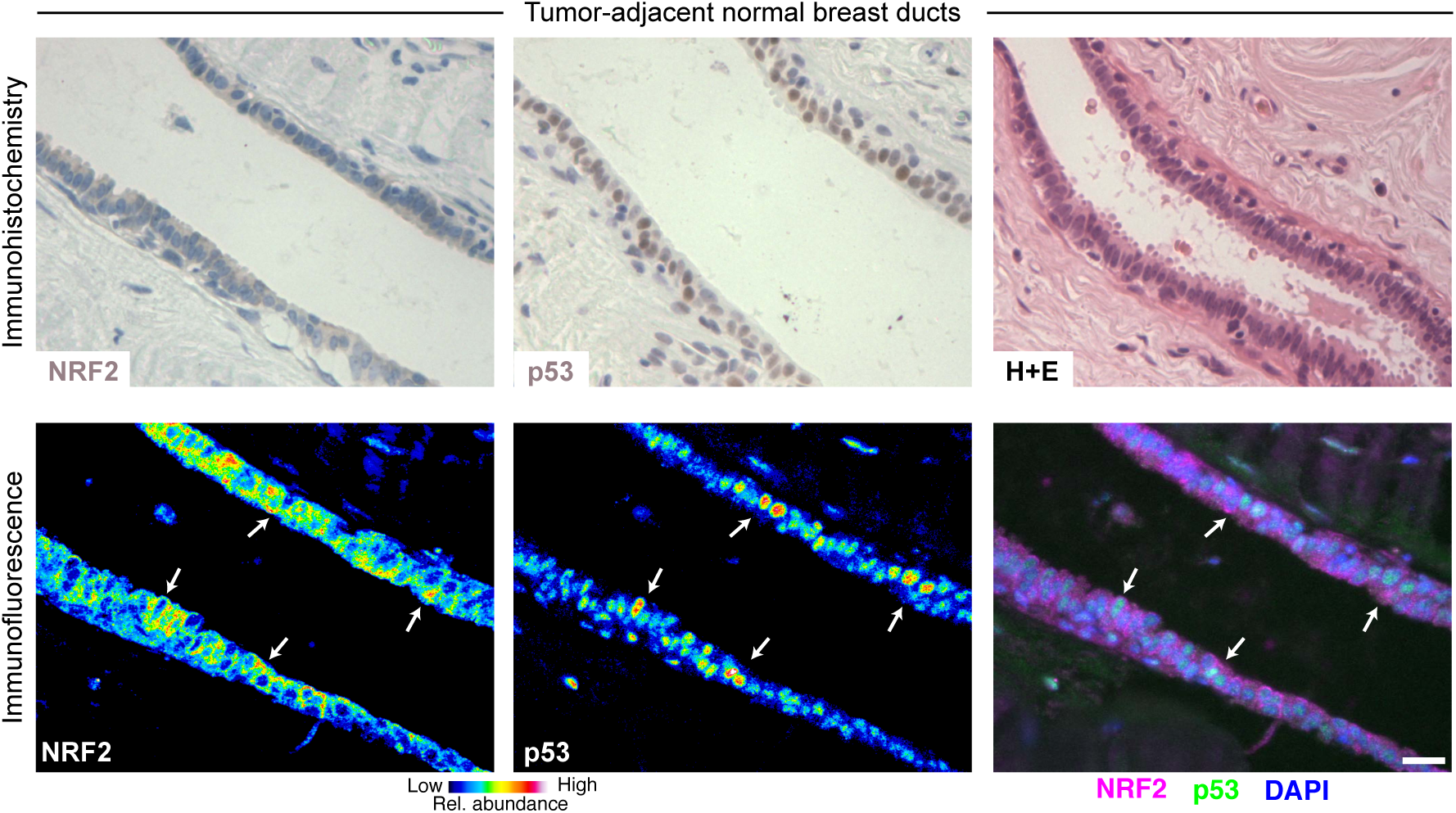
NRF2 and p53 are co-stabilized in breast epithelial ducts. Immunohistochemistry (upper) and immunofluorescence (lower) for NRF2 and p53 in tumor-adjacent normal breast ducts. Hematoxylin and eosin (H+E, upper right) histology is from a serial paraffin section for p53. Immunofluorescence is shown as representative pseudocolored images for NRF2 (left) and p53 (middle) are shown merged with DAPI nuclear counterstain (right). White arrows indicate concurrent NRF2 and p53 stabilization. Scale bar is 20 µm.

**Fig. S14.**
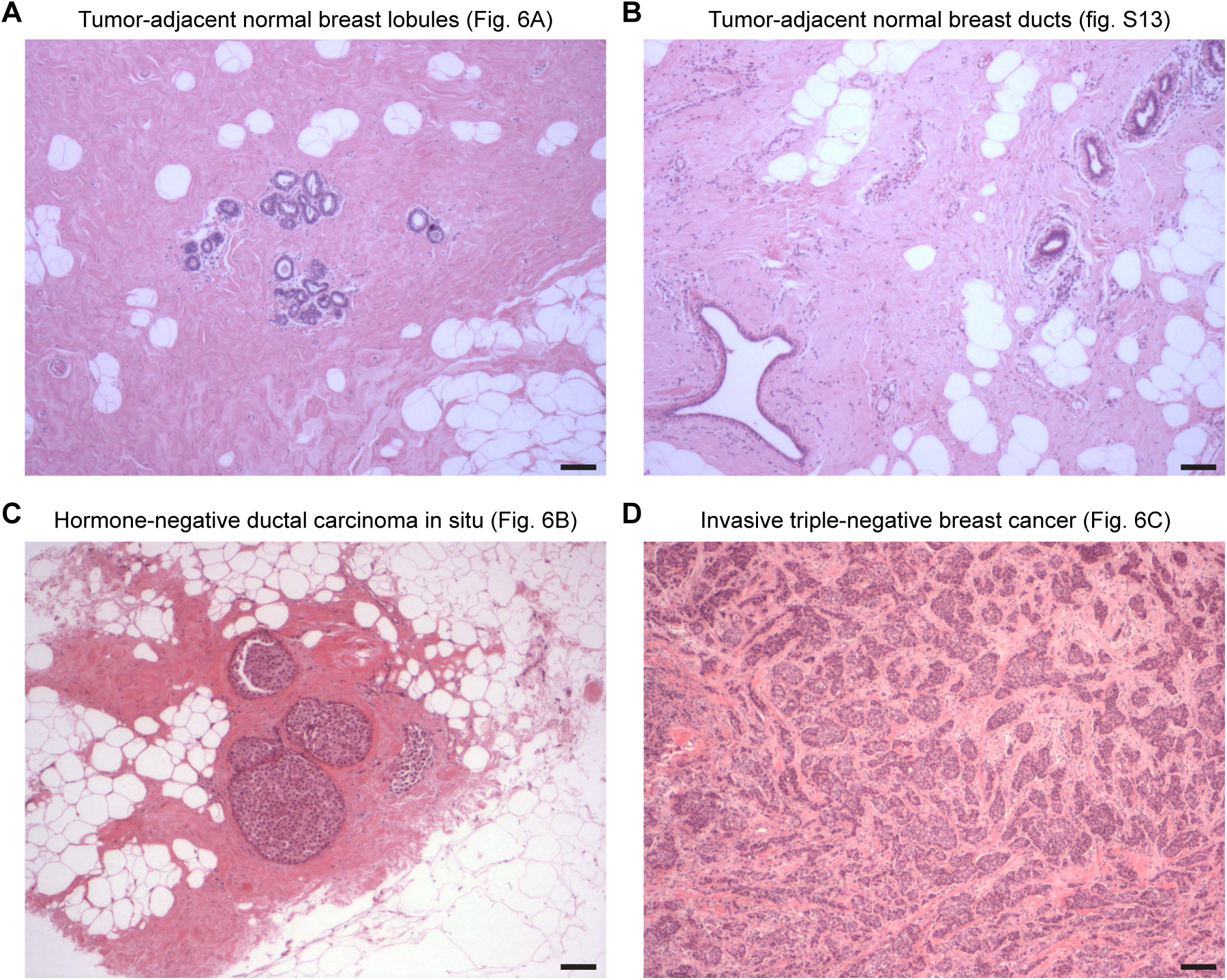
Low-magnification hematoxylin-eosin images of the tissues and tumors in the work. (**A**) Tumor-adjacent normal breast lobules from the specimen shown in Fig. 6A. (**B**) Tumor-adjacent normal breast ducts from the specimen shown in fig. S13. (**C**) Hormone-negative ductal carcinoma in situ from the specimen shown in Fig. 6B. (**D**) Invasive triple-negative breast cancer from the specimen shown in Fig. 6C. Scale bars are 200 µm.

**Table S1.**
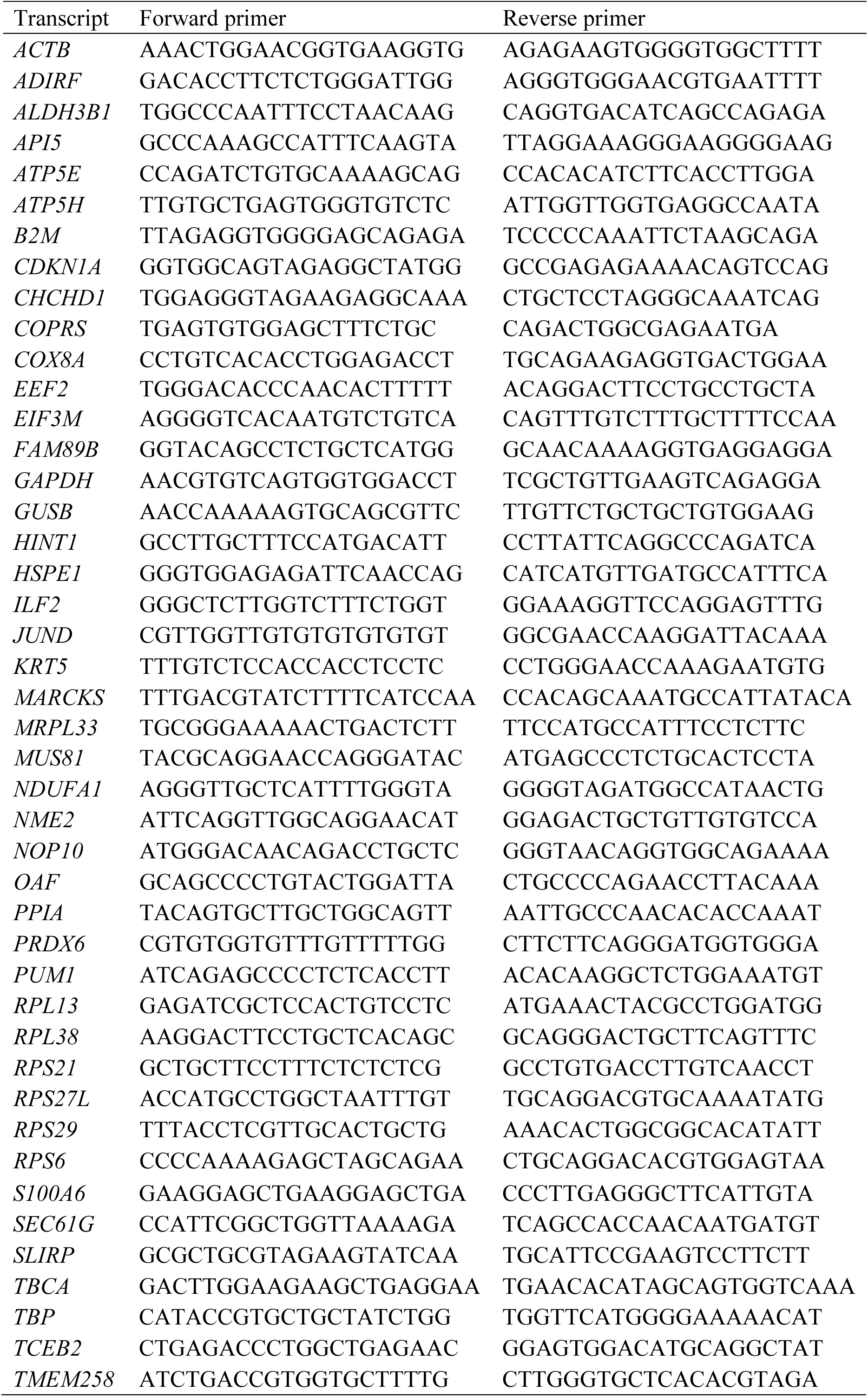
Quantitative PCR primer sequences.

**Table S2.**
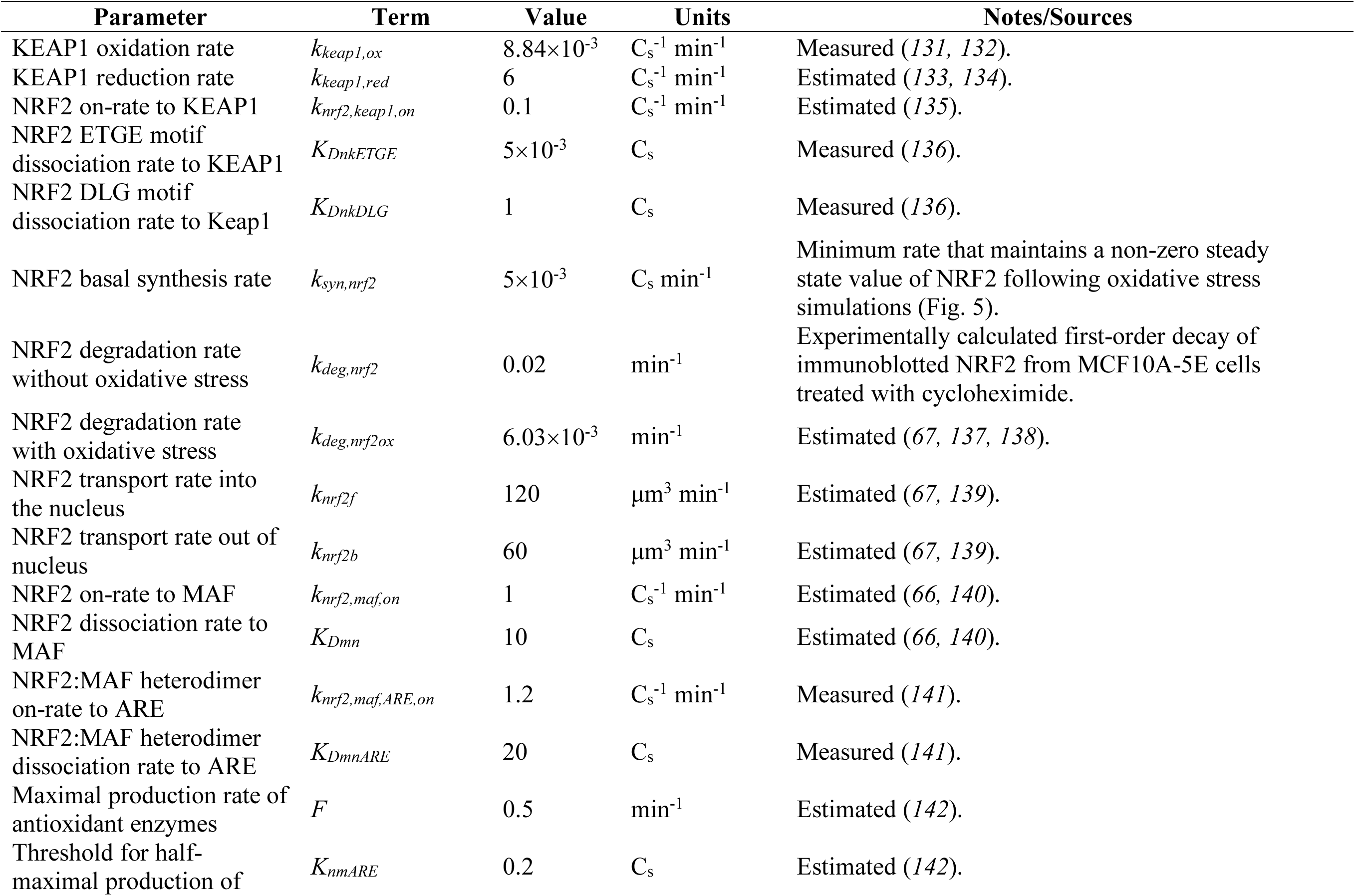
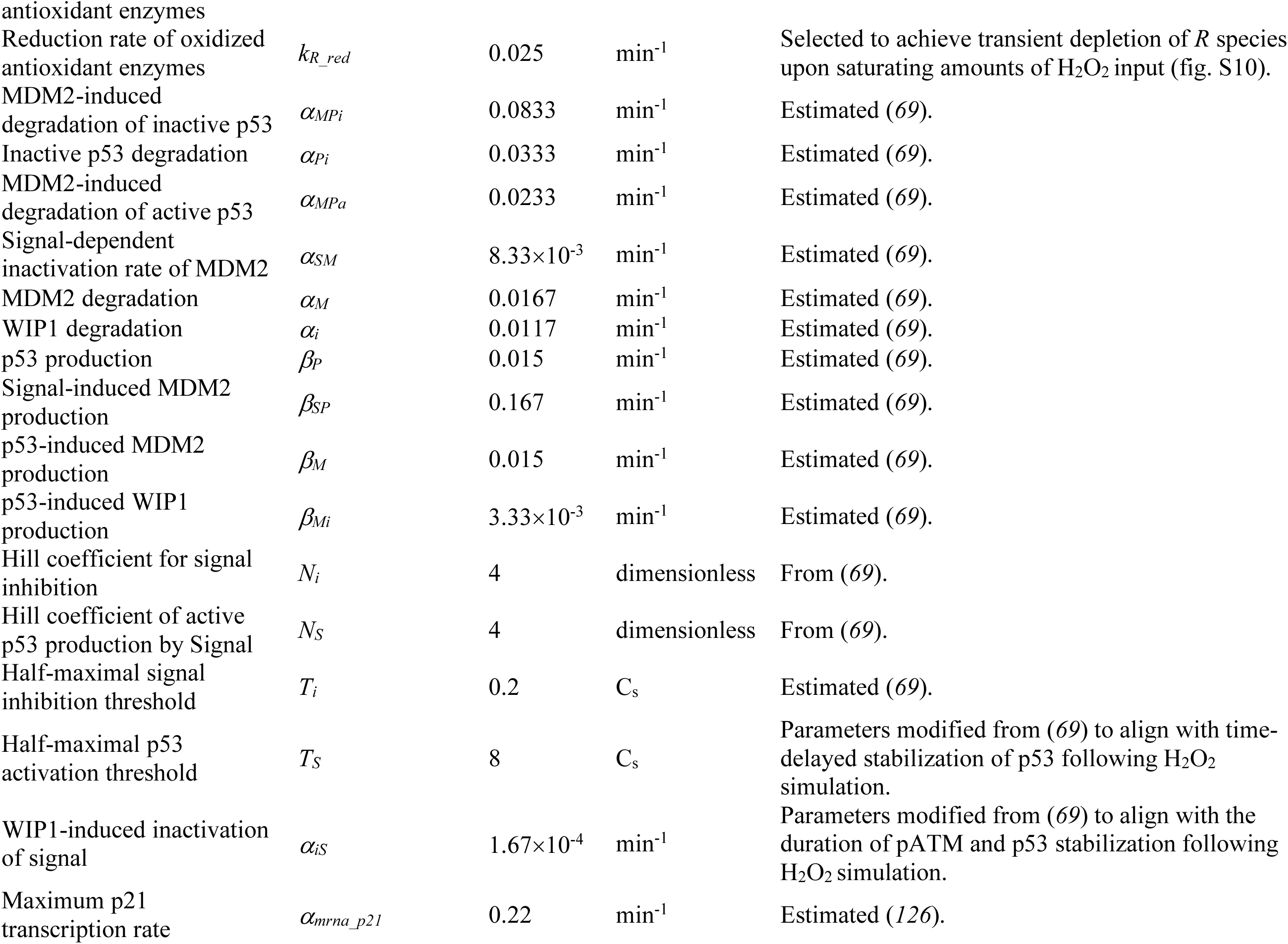
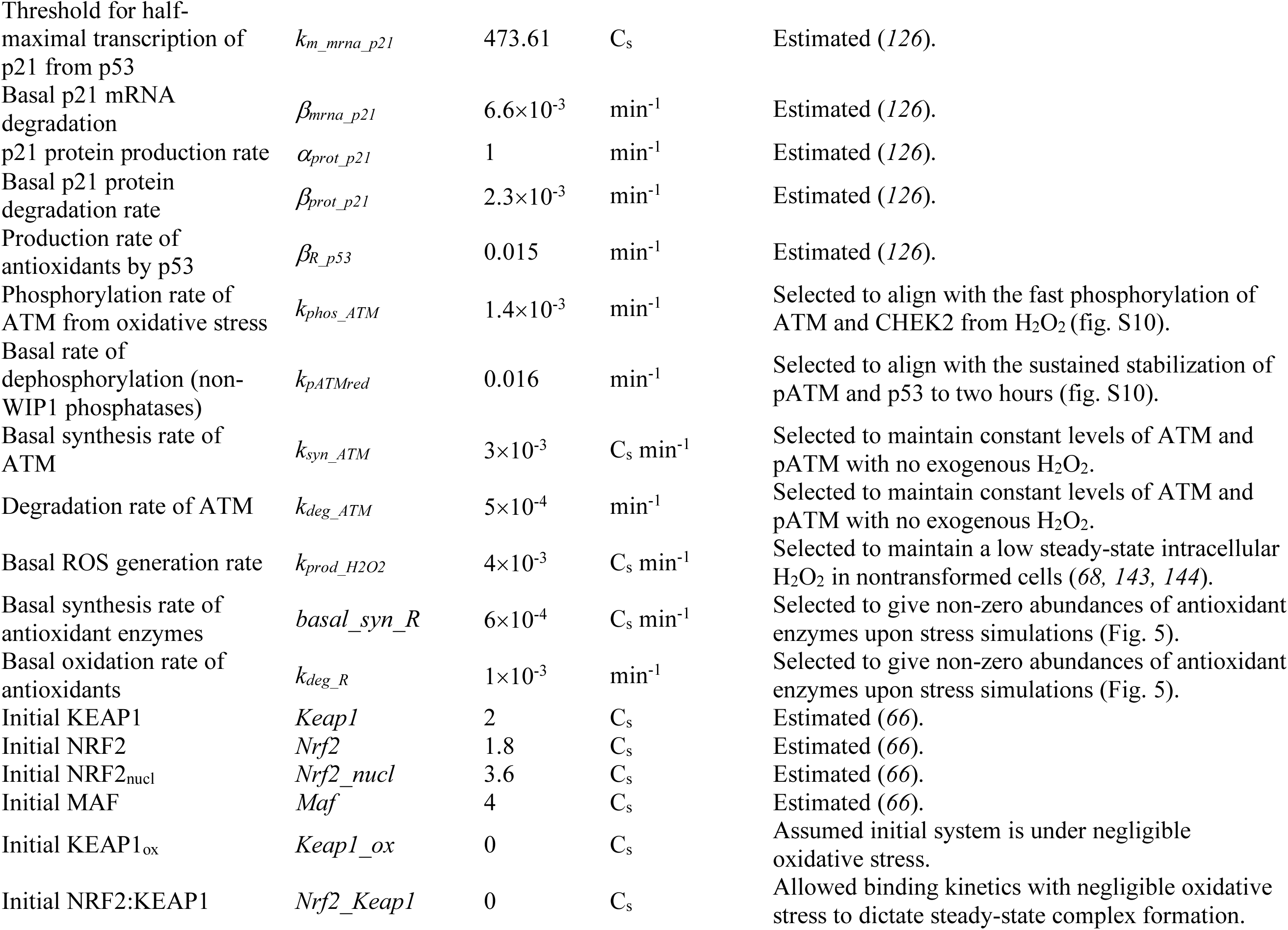
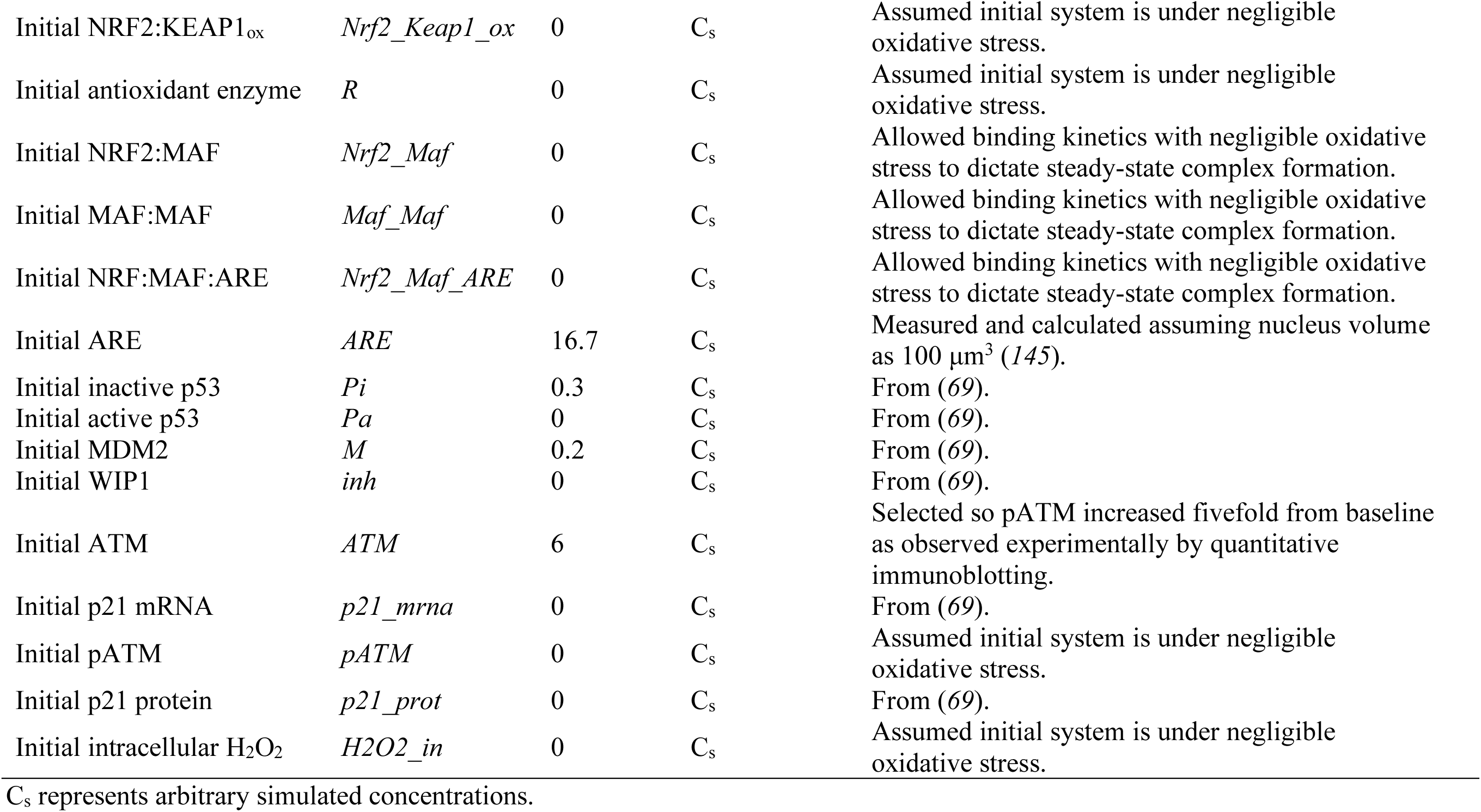
Parameter summary for the integrated NRF2–p53 computational model.

## Supplementary data files

**File S1. Gene Ontology enrichment analysis.** The Excel file contains the complete list of GO enrichments for transcripts in the 10-cell RNA sequencing dataset covarying with the median NRF2-associated signature (Spearman *ρ* > 0.5, *q* < 0.10).

**File S2. Gene set enrichment analysis of differentially abundant transcripts in MCF10A-5E and MCF10DCIS.com upon NRF2 knockdown compared to control.** The Excel file contains the complete list of gene sets significantly enriched in upregulated (sheet 1) and downregulated (sheet 2) genes upon NRF2 knockdown.

**File S3. NRF2–p53 computational model and associated files.** The Word file contains a URL to access the different computational models and parameter sets used in the work.

