## Supplementary material for "Sporadic activation of an oxidative stress-dependent NRF2–p53 signaling network in breast epithelial spheroids and premalignancies": File S3

**File S3. NRF2–p53 computational model and associated files.**

The .mat files specifying the steady state trajectories of the different models exceed the file upload limits of the journal. All files are available for download as a zipped archive here:

<https://virginia.box.com/s/hnkylh1rodn4t05ybgp2hv9ah39t7r83>
